# Phenotyping in the era of genomics: *MaTrics* – a digital character matrix to document mammalian phenotypic traits coded numerically

**DOI:** 10.1101/2021.01.17.426960

**Authors:** Clara Stefen, Franziska Wagner, Marika Asztalos, Peter Giere, Peter Grobe, Michael Hiller, Rebecca Hofmann, Maria Jähde, Ulla Lächele, Thomas Lehmann, Sylvia Ortmann, Benjamin Peters, Irina Ruf, Christian Schiffmann, Nadja Thier, Gabi Unterhitzenberger, Lars Vogt, Matthias Rudolf, Peggy Wehner, Heiko Stuckas

## Abstract

A new and uniquely structured matrix of mammalian phenotypes, *MaTrics* (*<u>Ma</u>mmalian <u>Tr</u>aits for Comparative Genom<u>ics</u>*) is presented in a digital form. By focussing on mammalian species for which genome assemblies are available, *MaTrics* provides an interface between mammalogy and comparative genomics.

*MaTrics* was developed as part of a project to link phenotypic differences between mammals to differences in their genomes using *Forward Genomics.* Apart from genomes this approach requires information on homologous phenotypes that are numerically encoded (presence-absence; multistate character coding*) in a matrix. *MaTrics* provides these data, links them to at least one reference (e.g., literature, photographs, histological sections, CT-scans, or museum specimens) and makes them available in a machine actionable NEXUS-format. By making the data computer readable, *MatTrics* opens a new way for digitizing collections. Currently, *MaTrics* covers 147 mammalian species and includes 207 characters referring to structure, morphology, physiology, ecology and ethology. Researching these traits revealed substantial knowledge gaps, highlighting the need for substantial phenotyping efforts in the genomic era. Using the trait information documented in *MaTrics*, previous Forward Genomics screens identified changes in genes that are associated with various phenotypes, ranging from fully-aquatic lifestyle to dietary specializations. These results motivate the continuous expansion of phenotype information, both by filling research gaps or by adding additional taxa and traits. *MaTrics* is digitally available online within the data repository Morph·D·Base (www.morphdbase.de).

## Introduction

### Background

Knowing and understanding the organisms around us has always been important for mankind and thus describing and comparing phenotypes has a long tradition that goes beyond the emergence of academic disciplines (e.g., Pruvost et al. 2011). The phenotype of an organism refers to its observable constituents, properties, and relations that can be considered to result from the interaction of the organism’s genotype with itself and its environment and include the anatomical organization of an organism, its physical properties, behaviour, ecological features, and lifestyle traits. They characterize an organism and contribute to biodiversity. Morphological* and anatomical* data usually make up the largest part of the phenotypic data available for a species. In mammalogy, specific skeletal, dental as well as body plan, visceral and physiological traits are traditionally used for differentiating species and for describing their inter- and intraspecific variability. Depending on preservation, this can also be applied to fossil species.

Advances in molecular biology and genetics over the last decades identified many genes and molecular mechanisms that are required for the development of many traits. This work relied primarily on studying model organisms such as the fruit fly (*Drosophila melanogaster*), the zebra fish (*Danio rerio*) or the mouse (*Mus musculus*). These models provided decisive insights into the genes behind basic developmental processes, including organ function and morphogenesis (Meunier 2012). Comparing developmental processes from model to non-model organisms opened the field for evolutionary developmental biology and explained the molecular basis of processes such as body plan evolution. However, there are some limitations on what model organisms can tell (Bolker 2012). For instance, insights from experiments on model organisms are restricted to the phenotypes present in that particular species. For example, rodents such as mice do not have canine teeth, making mouse an inappropriate model to study the molecular mechanisms required to make canine teeth. Furthermore, even if model organism research would reveal all genes that are associated with a given phenotype (e.g., the digestive system), it remains unknown which of these genes were altered during evolution and contributed to phenotypic changes between related species (e.g., mammals that specialized on particular diets).

With the development of sequencing technologies, sequencing and assembly of whole genomes became possible; the first was published in 1995 (of the bacteria *Haemophilus influenzae*, Fleischmann et al. 1995) and the mouse genome “only” was published in 2002 (Waterston et al. 2002). Due to advancements in high-throughput DNA sequencing, there are an increasing number of species for which sequenced nuclear genomes are available (e.g., Genome 10K Community of Scientists 2009; Teeling et al. 2018; Feng et al. 2020; Zoonomia Consortium 2020). This wealth of genomes provides a basis for comparative genomics* (“defined as the comparison of biological information derived from whole-genome sequences” and as discipline / methodology thus only started in 1995 (de Crécy-Lagard and Hanson 2018)). While comparative genomics often aims at identifying genomic elements that are conserved across species and thus likely have an evolutionarily important function (Nobrega and Pennacchio, 2004), comparative genomics can also be used to detect differences in functional genomic elements and associate them with phenotypic differences of species. For example, targeted analyses of genes associated with the formation of dentin (DSPP) and enamel (AMTN, AMBN, ENAM, AMELX, MMP20) across Mammalia and Sauropsida (including Aves, Crocodylia, Testudines, Squamata) showed an association between the loss of these genes and the loss of teeth (Meredith et al. 2009, 2014). Another example are losses of chitinase genes (CHIAs), enzymes that digest chitin, which preferentially occurred in mammalian species that have non-insectivorous diets (Emerling 2018).

Recent advances in comparative genomics follow the idea that convergent phenotypic evolution can be associated with convergent genomic changes e.g., gene loss (Lamichhaney et al., 2019). This assumption is one conceptual foundation of the general *Forward Genomics* approach that performs an unbiased screen for genomic changes that are associated with convergent phenotypic traits (Hiller et al. 2012; Prudent et al. 2016). This approach employs phenotype matrices and genome alignments to search for associations between convergent phenotypic traits and genomic signatures. These genomics signatures (e.g., candidate genes) need to be subjected to functional analyses to explain their association with the phenotype of interest. *Forward Genomics* identified many new links between genomic changes in genes as well as regulatory elements and various phenotypic changes such as adaptations to fully-aquatic lifestyles in cetaceans and manatees (Sharma et al. 2018a), echolocation in bats and toothed whales (Lee et al., 2018), reductions and losses of the mammalian vomeronasal system (Hecker et al., 2019a), the evolution of body armour in pangolins and armadillos (Sharma et al. 2018a), the absence of testicular descent (Sharma et al. 2018b), and the reduction of eye sight in subterranean mammals (Roscito et al., 2018; Langer and Hiller 2018).

### Development of MaTrics

A key requirement Forward Genomics is comprehensive and digitally-available phenotypic knowledge of species considered in the genomic screen. However, in contrast to genomic data, phenotypic data are not readily available in such a digitized form that it can be used by computer programs, not even for well-characterized species such as mammals with sequenced genomes. Research in Zoology and related fields assembled a rich body of phenotypic knowledge. But the information assembled over centuries is usually documented using natural language and thus in the form of texts unstructured for computer-programs and so the information is not machine-actionable* (Vogt et al. 2010). Whereas this is what researchers in Zoology and related fields used, and still use effectively, it limits research in other disciplines where substantial time investments are required to search and extract relevant phenotypic data from published descriptions. As a result, this important cultural and scientific heritage is underutilized in genomics and some disciplines of biomedical research.

Here, we address the need for digitally-available trait information by creating a phenotypic character matrix. Since the genome “encodes” all traits that have a genetic basis, genomes of many related species (such as mammals) enable Forward Genomic screens for many different traits with convergent changes. Thus, comprehensive information of many traits can be stored in a matrix form, where rows represent species and columns represent traits.

Constructing a comprehensive phenotypic matrix poses several challenges. While “simple” phenotypes that can be compiled relatively easily across several mammals, more complex phenotypes require experienced researchers in morphology, anatomy, physiology, veterinary science or related fields, since interpreting the collected information on phenotypes requires specialized knowledge on the terminology and taxon of interest. For example, the exact meaning of specialized terms might depend on the described taxon, the author, and the time of publication. Additionally, some terms might refer to spatio-structural properties, others to common function or presumed common evolutionary origin, or to a mixture of both. All this is well understandable to the experts, but, difficult for non-experts and even more so for computer algorithms. Thus, integrating the information on phenotypes in machine actionable form with other sources of data becomes exceedingly challenging and time-consuming (Lamichhaney et al. 2019; Vogt 2019). For integrative research a way is sought to exploit that knowledge without involving experts in each project.

To enable simpler use (and exploitation) of expert knowledge, more and more information is being digitized, stored, and made accessible online such as current journals or even older and classic books (Biodiverstiy Heritage Library). There are several databases for storing, editing and publishing information on phenotypes (mainly on morphological ones) covering various taxa (Table 1). All of them have their own purpose and relevance, but none of them so far fulfils the requirements of *Forward Genomics* and other efforts to link phenotypic to genomic differences; most neither provide information on the same character across the selected species nor is the information numerically coded to be directly useful to other computer analysing programs. However, tables on morphological traits for phylogenetic analyses fulfil these requirements, but do not focus on extant mammals with sequenced genomes. Furthermore, while inference of homologies in genomic data (nucleotide sequence alignment) is fully automated, homology analyses (character alignment) of phenotypic data cannot be executed by computer algorithms so far. This is irrespective of the type of basic information available (digitized literature, 2D/3D scans of museum specimens). Matrices usable to link phenotypic differences between species to genomic loci first need to provide homology information.

**Table 1.** Examples of data repositories in which phenotypic data of different vertebrate taxa are collected. The table lists the projects with their URL and aim and/or type of information that is stored and, if available, references in which the project is introduced.

| Project | Link | Aim, type of information | Reference |
| --- | --- | --- | --- |
| Morphobank | http://www.morphobank.org | Homology of phenotypes over the web; building the Tree of Life with phenotypes, publicly accessible containing images and matrices | O'Leary and Kaufmann 2011 |
| Digimorph | http://www.digimorph.org | A National Science Foundation Digital Library at The University of Texas Austin, a dynamic archive that holds high-resolution X-ray computed tomography of biological specimens |  |
| Morphbank | http://www.morphbank.net | A continuously growing database of Biological Imaging and stores images that scientists use for international collaboration, research and education |  |
| Morphological Image database | http://people.pwf.cam.ac.uk/rja58/database/morphsite_bmc07.html |  | Asher 2007 |
| Phenoscape | http://kb.phenoscape.org | Data resource that is ontology-driven and contains information about mutant zebrafish (*Danio rerio*) phenotypes curated by the zebrafish model organism database, ZFIN at http://zfin.org | Ruzicka et al. 2015; Edmunds et al. 2015 |
| TOFF | http://toff-project.univ-lorraine.fr | An open source repository focusing on fish functional traits. It aims to combine behavioural, morphological, phenological, and physiological traits with environmental measurements | Lecocq et al. 2019 |

To enable the full use of *Forward Genomics*, a trait matrix of mammalian phenotypes was developed that fulfils the above-mentioned criteria. Here, we introduce *MaTrics* (*<u>Ma</u>*mmalian *<u>Tr</u>*aits for Comparative genom*<u>ics</u>*), the first and newly established matrix providing information on phenotypic traits of mammals.

## Design and coding* principles of *MaTrics*

*MaTrics*, version 1.0, release January 2021 (in the following still referred to as *MaTrics* only) is implemented in the online data repository* Morph·D·Base (www.morphdbase.de, Grobe and Vogt 2009) and publicly available.

### Principles and data entry

*MaTrics* meets all requirements of *Forward Genomics*. a We primarily focused on mammalian species for which genome sequences are available. Some basic principles of *MaTrics* are described herein, a detailed user’s guide is available online (Wagner et al. 2020). Different types of phenotypic traits were considered (see below) and in each case homology was assured.

In case a phenotypic trait has several different expressions, it must be coded as a multistate character. According Sereno (2007), the *character* part in a multistate character comprises not only the *locator* but additionally a *variable* (V – the aspect that varies) and a *variable qualifier* (q – the variable modifier). The *character states* of a multistate character in *MaTrics* are numerically coded by 2 to n (Fig. 1B, Table 2). For example, the height of the mandibular canine teeth in relation to the level of the occlusal height (averaged) of the cheek teeth are coded as *short* (2), *occlusal height* (3) or *long* (4) (Fig. 1B).

**Fig. 1.**
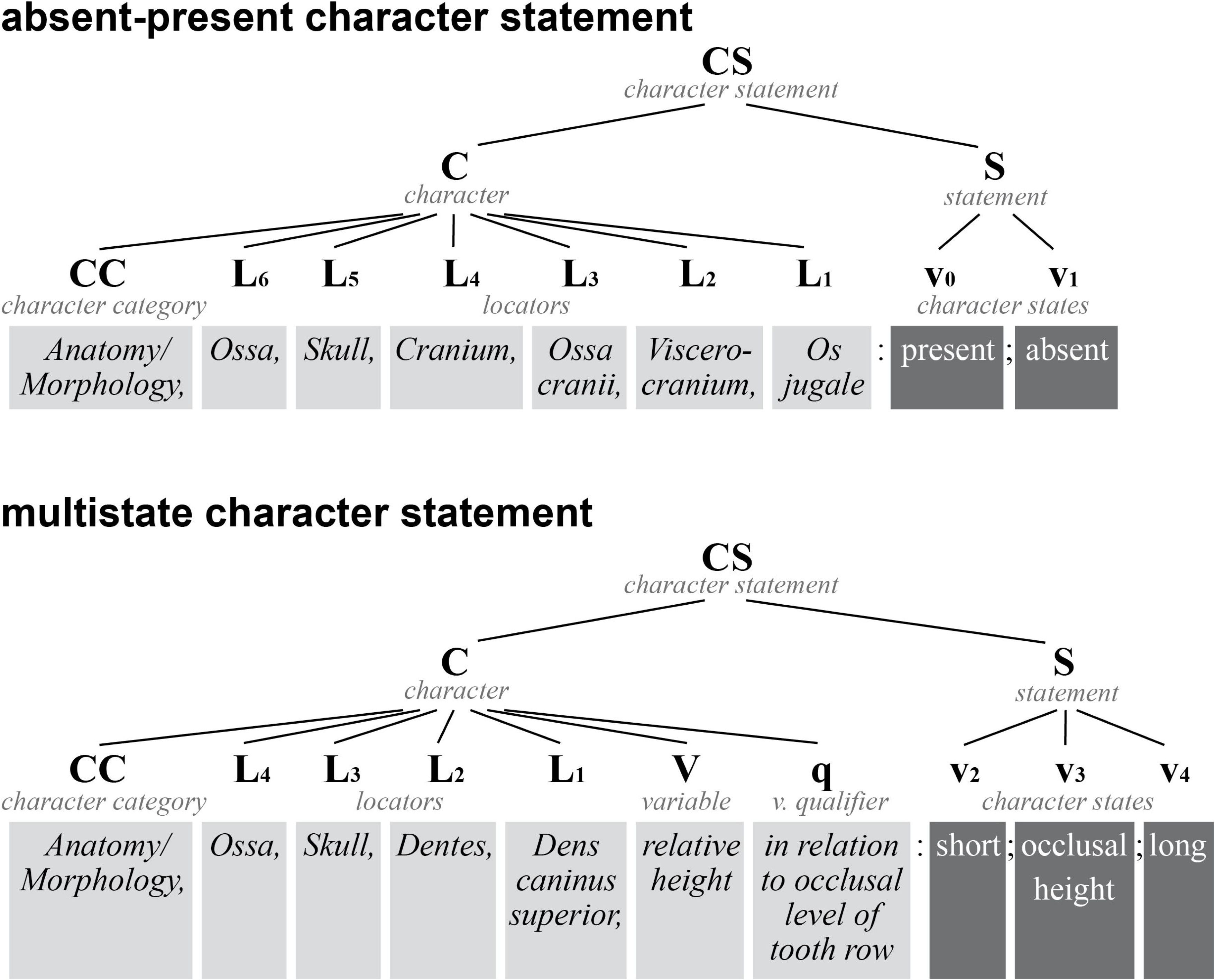
Schematic illustration showing how phenotypic traits are reflected in *character statements* and in the character labels in *MaTrics*. The basic nomenclature is based on Sereno (2007: table 4, Scheme 3), see A1 and B1. A) Illustrates the structure for characters which can be described with only two *character states*: absent and present. B) Illustrates the structure for characters which require more than two *character states* (multistate characters). A2 and B2 give the terminology for the examples from *MaTrics* named in A3/B3. Sereno’s (2007) terminology recognizes *character statements* (CS) consisting of *characters* (C) and *statements* (S). The *character* is represented by a (list of) *locators* (L_n_,…L_1_; hierarchically organized and forming the structure tree) and optionally the *variable* (V) and the *variable qualifier* (q). The different expressions of the *variable* are given as *character states* (v_0_,… to v_n_) representing the *statement*. A4/B4 are examples how this nomenclature is given in the character label in *MaTrics*. The *character states* are defined in the “states” field and assigned to each cell of *MaTrics*. Whether a character can be described by the two states absent/present or several states is indicated in the character label by the addition [a/p] and [m], respectively.

**Table 2.**
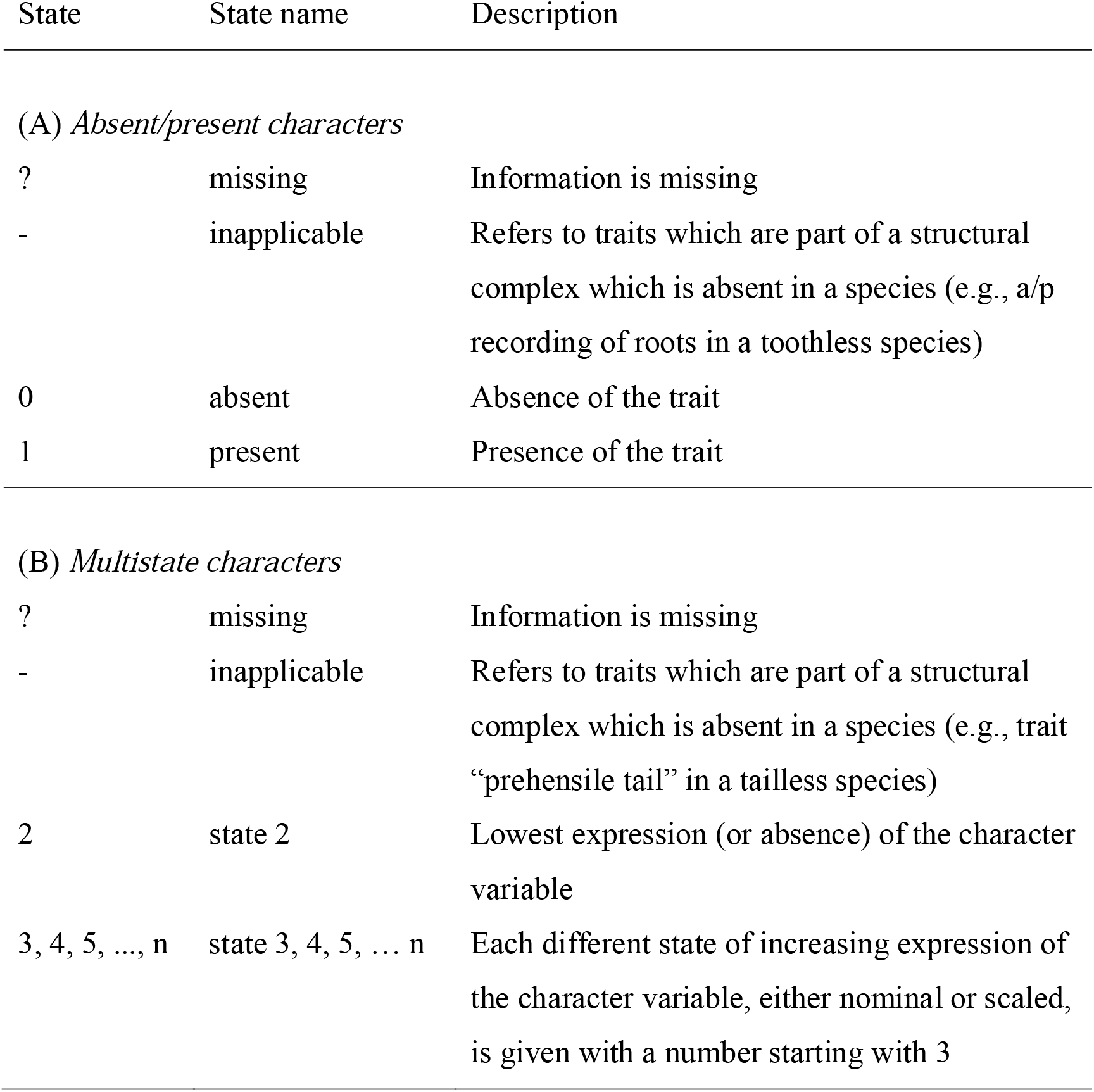
The numerical coding options for (A) absent/present and (B) multistate characters in *MaTrics*. The numerals 0 and 1 refer to the *character states* ‘absent’ and ‘present’, thus, the coding for multistate character states starts at 2 and continues to

According to Sereno (2007), a (phenotypic) trait of an operational taxonomic unit (OUT; here the selected mammalian species) can be represented in a *character statement* that is composed of two parts: *character* and *statement*, and can be divided into four types of logical components (Sereno 2007:Table 4): one or more *locators*, a *variable*, and a *variable qualifier* as parts of the character and a *character state* as the *statement.* Not all these components are needed in any case, but a *locator* and a *character state* are the minimum (representing *character* and *statement*). Thus, each *character* consists of at least one *locator* (L – the morphological structure, the structure bearing the trait) and the *statement* of the *character state* (v – mutually exclusive condition of a character) (Fig. 1). Specifying a *locator* and a *character state* is sufficient in case of absent-present *character statements\** (numerically coded by 0/1; Fig. 1A, Table 2). Following Sereno’s (2007) coding scheme, each character in *MaTrics* is named with a label starting with a single *locator* or a sequence of *locators* starting with L_n_ to L_1_ (the trait bearing structure), which provide all information necessary for unambiguously identifying and locating the trait within the OTU. The sequence of locators (L_n_ to L_1_ as illustrated in Fig. 1) in the character label is hierarchically organized. While Sereno (2007) developed his coding scheme primarily for structural traits, we extended it here and applied it also to ecological or behavioural traits.

In case a phenotypic trait has several different expressions or patterns, it must be coded as a multistate character. According Sereno (2007), the *character* part in a multistate character comprises not only the *locator* but additionally a *variable* (V – the aspect that varies) and a *variable qualifier* (q – the variable qualifier). The *character states* of a multistate character in *MaTrics* are numerically coded by 2 to n (Fig. 1B, Table 2). For example, the height of the mandibular canine teeth in relation to the level of the occlusal height (averaged) of the cheek teeth are coded as *short* (2), *occlusal height* (3) or *long* (4) (Fig. 1B).

A key consideration when generating *MaTrics* was to clearly document the source(s) for each phenotypic entry. In *MaTrics*, the *character* part of each *character statement* therefore possesses a short textual definition that is taken from published sources (journals, text books, online references) and includes references to relevant ontology terms from various biomedical ontologies (the following online resources were used for identifying adequate terms: Ontology Lookup Service, OLS, https://www.ebi.ac.uk/ols/index, Jupp et al. (2015); Ontobee, https://www.ontobee.org, Xiang et al. (2011); Bioportal, https://bioportal.bioontology.org, Musen et al. (2012)). If no adequate definition was available, we provided our own definitions and clearly marked them as such.

The dimensions of *MaTrics* are defined by the number of rows (OTUs) and columns (characters) that result in a specific number of cells (rows × columns). These cells primarily contain the character states. Morph·D·Base enables the addition of further information such as references, photos, illustrations, or museum specimen IDs to each matrix cell. All character states recorded, thus each cell of *MaTrics* is linked to at least one supporting reference. This refers either to citations from the literature (e.g., published journal articles, books, reliable scientific online resources) or to primary data sources. These data sources can cover IDs of museum specimens or direct links to media (e.g., photographs; microscopic and electron microscopic (TEM and SEM) images, magnetic resonance (MRI), computed tomography (, μCT), or even synchrotron data) which are directly uploaded in MDB. As a result, researchers using *MaTrics* can trace the information to at least one original source.

Phenotypic traits coded in *MaTrics* represent by default adult states. Fetal structures or traits that belong to perinatal or not yet fully-grown stages are explicitly indicated as “fetal” (fetal is used as *locator* L_n_ in the character label). Traits referring to other ontogenetic stages can be coded in a similar way.

Phenotypic traits included in *MaTrics* represent by default adult stages. Fetal structures, or traits that belong to perinatal or not yet fully-grown stages are explicitly indicated by placing “fetal” in front of the *locator* L_n_ in the character label. Traits referring to other ontogenetic stages could be considered in a similar way.

The *MaTrics* or individual characters can be exported as a Nexus* file that provides data in a structured way and can be used as input in various software analysis tools.

### Specificities of MaTrics

The primary motivation to generating *MaTrics* was to create a research tool for linking phenotypic differences between species to differences in their genomes. This is the main reason why intraspecific variations of traits such as sexual dimorphism were not considered. Another specificity is that *character states* (presence/absence; multistate) do not encode character polarity. Researchers can decide for each project individually whether to use and determine polarity or not. The characters might be further analysed (e.g., polarity analyses using out-group comparison) if considered for phylogenetic studies or gene loss analyses. Finally, character dependencies were not specifically accounted for during the choice and coding of traits. For each research question, specific characters of interest were added to *MaTrics.* Similarly, for different projects, characters can be selected individually to be retrieved from *MaTrics* for other use. Character dependencies can be avoided or reduced in this way, if needed.

## Current status:*MaTrics*

To date, *MaTrics* contains 207 characters for 147 mammalian species to date, resulting in a total of 30,282 documented character states. 153 of the 207 characters (74%) are described as absent-present characters and the remaining 53 (26%) are multistate characters. The mammalian species considered in *MaTrics* include two representatives of Monotremata, five of Marsupialia and 140 of placental mammals (supplementary material Table S1). The number of species from each order neither represents the respective diversity nor the morphological disparity of mammalian orders, as the primary criterion for the inclusion in *MaTrics* was the availability and suitable quality of whole genomes. The characters in *MaTrics* cover structural, ecological, ethological, and physiological phenotypic traits (Table 3). All, but one character (*organum vomeronasale*), refer to the adult stage. For one character (*os jugale*), the recording is 100%, so all cells contain coded and referenced character states. Some traits were specifically included for the study in subsets of the listed mammals, and therefore the recording purposely is less complete (for coding status see Table S2).

**Table 3.** Gross categories of 206 characters included in *MaTrics* and number of characters in these categories

| Gross category | Subcategory | n |
| --- | --- | --- |
| Anatomy/Morphology |  | 164 |
|  | Body plan | 1 |
|  | Cranial skeleton | 30 |
|  | Dentition | 94 |
|  | Gastrointestinal tract | 5 |
|  | Head | 4 |
|  | Integument | 3 |
|  | Postcranial skeleton | 26 |
|  | Sense organs | 1 |
| Ecology |  | 30 |
| Ethology |  | 5 |
| Physiology |  | 6 |
| Embryonic |  | 1 |
| Total |  | 206 |

## Notes on application

The primary motivation for creating *MaTrics* was to provide fully referenced phenotypic information for applications in comparative genomics, especially the *Forward Genomics* approach. The creation and filling of *MaTrics* and studies applying *Forward Genomics* were developed in parallel within the mentioned project. So, phenotypes were coded in *MaTrics* were partially successfully used in earlier studies and simpler shorter tables e.g. by Sharma et al. (2018a) who identified various convergent gene losses associated with some specific convergent mammalian phenotypes. They showed convincingly that tooth and enamel loss are associated with the loss of ACP4 (a gene that is associated with the enamel disorder amelogenesis imperfecta), and that the presence of scales is associated with the loss of the gene DDB2 (which detects substances resulting from UV-light and helps to induce DNA repair). The fully aquatic lifestyle is associated with the loss of MMP12, a gene associated with breathing adaptation. The documented loss of these genes in some mammalian species is functionally explainable either as a consequence of trait loss (the genes ACP4 and DDB2 have no function after trait loss) or as putative adaptive genomic alteration, causing novel phenotypes (MMP12-loss is associated with novel lung functions in aquatic mammals) (Sharma et al. 2018a). Such results might help to better understand some related human diseases, as for example in the case of DDB2 whose mutations cause xeroderma pigmentosum which manifests in hypersensitivity to sunlight (Rapic-Otric et al. 2003).

Another study investigated the gene losses associated with the reduction of the vomeronasal system (VNS) in several mammals. A genomic comparison of 115 mammalian genomes confirmed that *Trpc2* is an indicator for the functionality of the VNS (Hecker et al. 2019a). Moreover, it indicated a loss of functionality of the VNS in seals (Phocidae) and otters (Lutrinae). Morphological data is scarce for seals and there is no data for otters (Hecker et al. 2019a; Zhang and Nikaido 2020). A study to test the accuracy of the suggested predictability is under way. This study is an example for testing genotype-phenotype associations in non-model organisms and shows the potential of the combination of comparative morphological and genomic approaches.

However, the relevance of *MaTrics* is by no means restricted to the *Forward Genomics* approach. Characters were also included in *MaTric* for the usage in the contemporary study to explore evolutionary conditions associated with the loss of genes related to convergent evolution of herbivorous and carnivorous diet in mammals (Hecker et al. 2019b). This study included 52 placental species and suggests that the lipase inhibitor gene PNLIPRP1 is preferentially lost in herbivores, whereas the xenobiotic receptor NR1I3 is preferably lost in carnivores. Even though the authors put forward hypotheses, the lack of accessible data on mammalian diet preferences made it difficult to test whether gene losses are associated with dietary fat content and diet-related toxins. Investigating whether convergent gene loss is associated with similar dietary preferences may additionally hold information on whether gene losses might be adaptive (Albalat and Cañestro 2016). Consequently, an ongoing study records dietary categories in *MaTrics* that allow a semi-quantitative of dietary fat content (associated with PNLIPRP1) and diet-related toxins (associated with NR1I3) (Wagner et al. ####). This study provided evidence that the convergent loss of both genes is associated with the convergent evolutionary change of dietary preferences, i.e. the consumption of a diet with reduced fat and toxin contents. The hypotheses of Hecker et al. could be refined and also the evolutionary setting could be reconstructed.

Future analyses using *MaTrics* have the potential to test how gene losses and dietary composition are related to the presence/absence of structures or organs associated with digestive processes. Even further, it allows investigating whether evolutionary changes in diet composition are not only associated with the loss/presence of single molecules (e.g., lipase inhibitor, xenobiotic receptor), but also with changes in complex structures and their associated gene. For instance, it is interesting to note that first statistical investigations (methods given in document S3) have not yet proven a significant association between the presence of a gall bladder and the diet (p=0.74) as well as the lipase inhibitor gene PNLIPRP1 (p=0.49). This observation motivates the further development of *MaTrics*, i.e. by adding further traits and species.

These two studies show how genomic and morphological studies are entangled: current knowledge of morphology serves as basis for creating phenotypic trait matrices like *MaTrics* which – on the other hand – forms the basis of genomic research, especially the *Forward Genomics* approach. Hypotheses associated with findings of candidate loci, may in turn inspire further morphological research.

The most obvious application are morphological studies. Although mammal dentitions are well studied and a lot is known about teeth number, form, and shape in particular in relation to dietary specialization (see Thenius 1989; Hillson 2005; Ungar 2010), we still have many gaps of knowledge, e.g., concerning functional adaptations and evolutionary transformations. Thus, Sole and Ladevèze (2017) aimed to put forward new ideas on how the basic mammalian tribosphenic molar was transformed to sectorial teeth in hypercarnivorous mammals. They (Sole and Ladevèze 2017) included only carnivores as defined by flesh-eating and the presence of carnassial teeth, representatives of the living Carnivoramrpha (including the extinct Nimravidae) and Dasyuromorphia, as well as from the extinct Sparassoodonta, Oxyaenodonta, and Hyaenodontida in their study. Comparing the cusp pattern/morphology of the upper and lower molars of these species Sole and Ladevéze (2017:fig. 4) derived a scheme for the morphological evolution of the sectorial teeth in hypercarnivorous mammals. They also aimed at providing new arguments to discuss the developmental aspects of the evolution of hypercarnivory by associating their morphological observations with ontogenetic studies. The latter highlighted the importance of the expression of ectodysplasin A (Eda): increased levels are able to modify the number, shape, and position of cusps in mice during tooth development (Kangas et al. 2004). Further, Häärä et al. (2012:3189) showed – again in mice – that “Fgf20 is a major downstream effector of Eda and affects Eda-regulated characteristics of tooth morphogenesis, including the number, size, and shape of teeth. Fgf20 function is compensated for by other Fgfs”. Inspired by the observations and the model of Solé and Ladevèze (2017), we started a study with a subsample of Carnivora (Table S3) collected in *MaTrics* with two aims: firstly, to test the suitability of *MaTrics* in comparative morphological studies and, secondly to set the basis to proceed with genome wide searches for genomic causes correlated with the loss of cusps. This seems to be promising with the development of new methods to include searches for regulatory elements (see below).

For the selected Carnivora (Table S4) the absence and presence of individual tooth cusps for the fourth upper premolar (P^4^) and all molar teeth were recorded in *MaTrics*. The nomenclature of the cusps followed Thenius (1989, exemplified in Fig. 2). The detailed descriptions of cusp patterns for the species are given in the supplementary document S5 and examples are illustrated in Fig. 3 and detailed in Table S6. Some of our results confirmed the observations of Solé and Ladevèze (2017), who focused on carnivores as defined by the presence of carnassials. We confirm that parastyle and protocone of the P^4^ are generally reduced in hypercarnivorous carnivorans. Interestingly, both structures are more reduced in the Canidae and the polar bear (*Ursus maritimus*) than in the members of the Felidae and Hyaenidae. Solé and Ladevèze (2017) reported that in the upper molars protocone, paraconule and metaconule are reduced in hypercarnivorous mammals which is also in line with our findings.

**Fig. 2.**
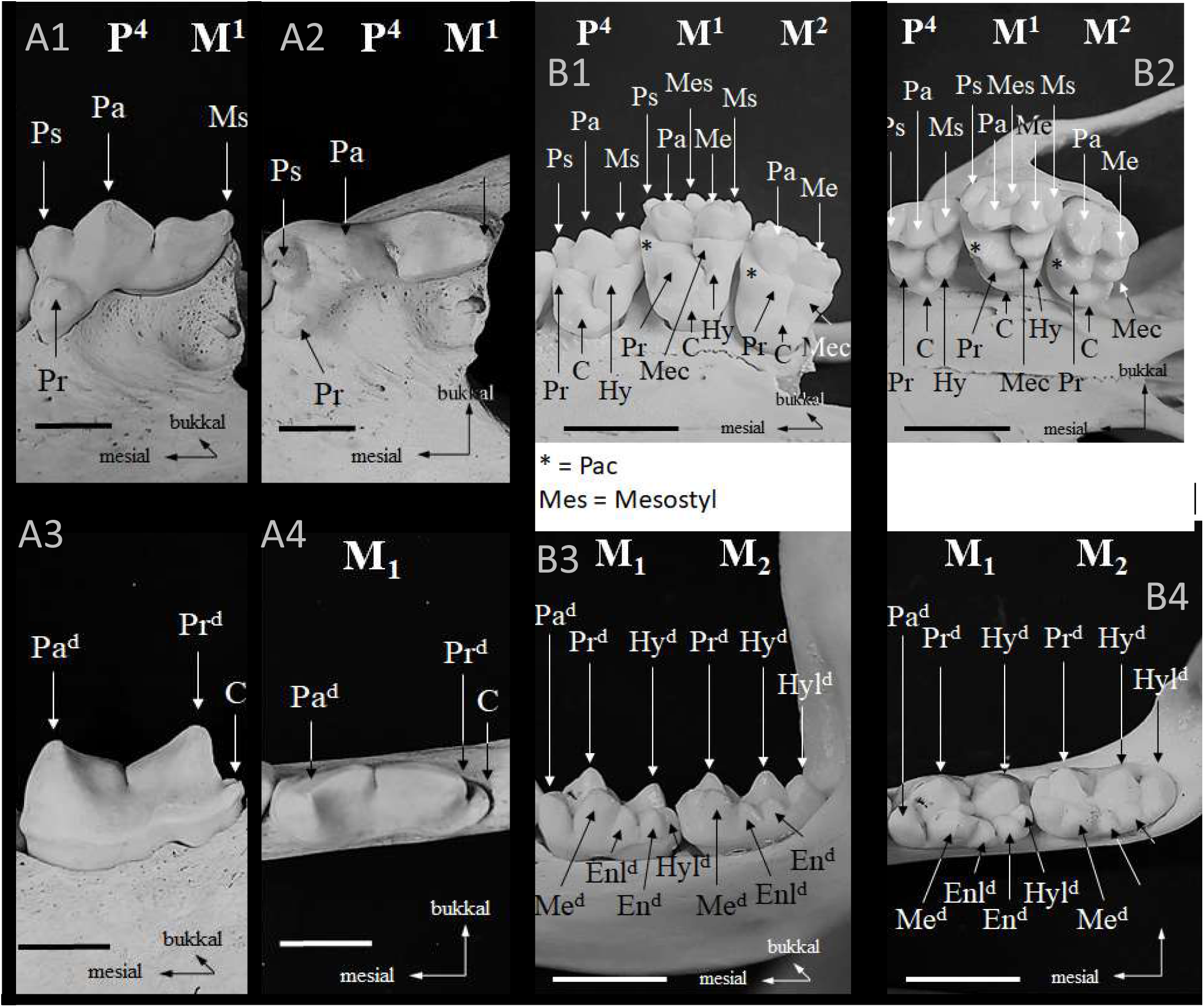
Some examples for the presence of cusps in the studied Carnivora. A) the spotted hyaen *Crocuta Crocuta* MTD B4936, B) the red panda *Ailurus fulgens* MTD B17478, C) the panda *Ailuropda melanoleuca* ZMB_Mam_17246 and D) the Weddell seal *Leptonychotes weddellii* MTD B5029. For each species the upper P4 and molars (1, 2) and lower molars (3, 4) are illustrated as present and the cusps labelled. The teeth are photographed in lateral (1, 3) and occlusal (2, 4) view. Abbreviations alphabetically: **En^d^** – entoconid, **Enl^d^** – entoconulid, **Hy** – hypocone, **Hy^d^** – hypoconid, **Hyl^d^** – hypoconulid, **Me** – metacone, **Mec** – metaconule, **Me^d^** – metaconid, **Mes** – mesostyle, **Ms** – metastyle, **Pa** – paracone, **Pac** – paraconule, **Pa^d^** – paraconid, **Pr** – protocone, **Pr^d^** – protoconid and **Ps** – parastyle

**Fig. 3.**
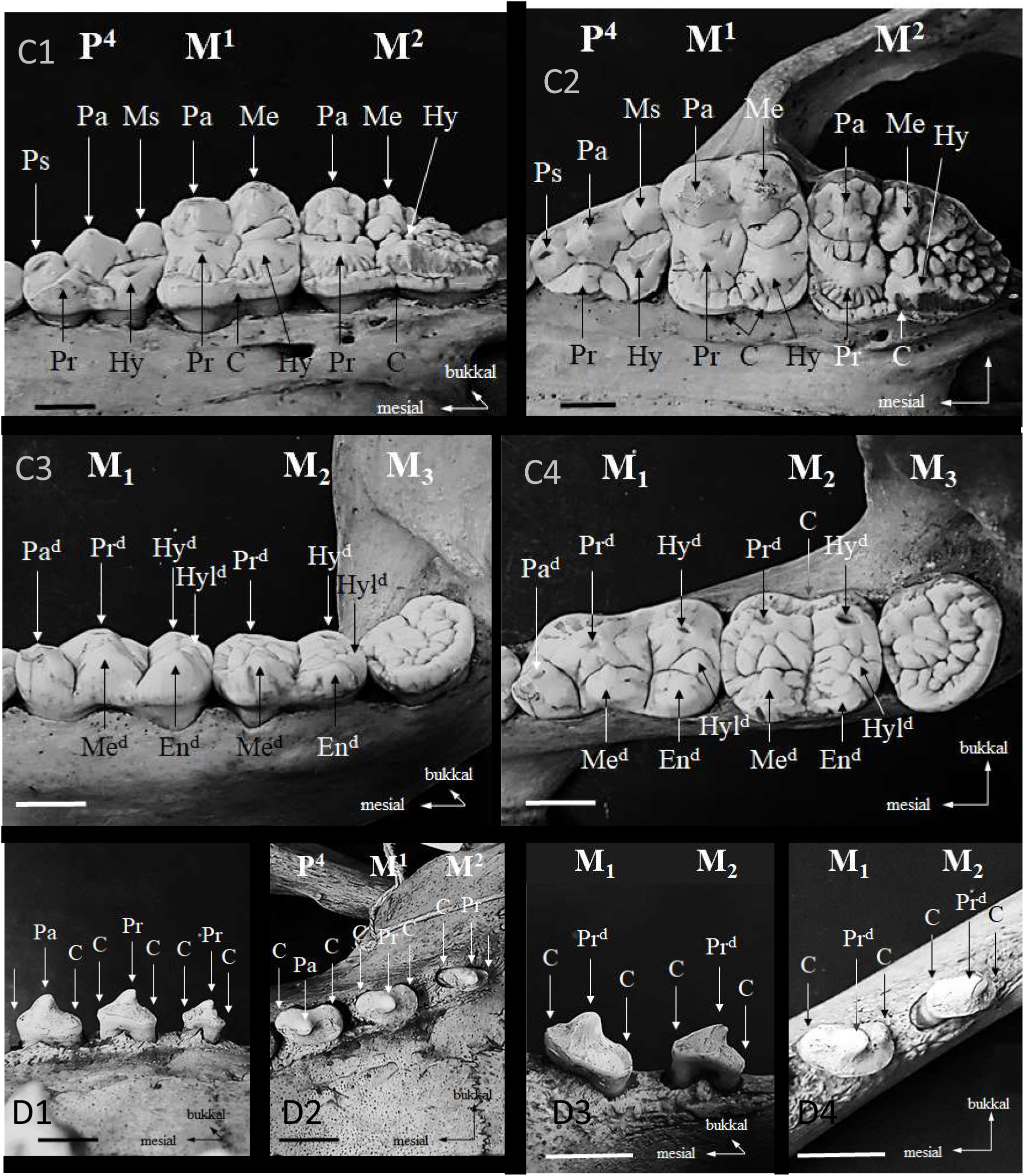
The presence and absence of cusps in P^4^ and M^1^ exemplified in A) the spotted hyaena *Crocuta Crocuta*, B) the red panda *Ailurus fulgens*, C) the panda *Ailuropda melanoleuca* and D) the Weddell seal *Leptonychotes weddellii* (teeth illustrated in Fig. 3). Abbreviations as they appear in table: **Ps** – parastyle, **Pa** – paracone, **Pr** – protocone, **Ms** – metastyle, **Hy** – hypocone, **Mes** – mesostyle, **Me** – metacone, **Mec** – metaconule, and **Pac** – paraconule

These structures are reduced in the Canidae, and totally absent in the Felidae and Hyaenidae. Solé and Ladevèze (2017) also found, that metaconid and talonid are generally lost in hypercarnivorous mammals, especially felid-like and hyaenid-like hypercarnivores. Based on our study, we found that metaconid and talonid are completely reduced only in the Felidae (except the cheetah, *Acinonyx jubatus*) and the spotted hyena (*Crocuta crocuta*). Like in the Canidae and the striped hyena (*Hyaena hyaena*), both structures are also present in *Ursus maritimus*. The specialized hypercarnivorous diet of several Feliformia lead to an extreme reduction of the tribosphenic molar, whereas the Canidae and *Ursus maritimus* also eat fruits and vegetables and therefore need crushing structures. The presence of protocone and talonid seems to be necessary for an omnivorous diet (Solé and Ladevèze 2017), but based on our study we can confirm that this is also true for herbivorous species (e.g., red panda, *Ailurus fulgens*; giant panda, *Ailuropoda melanoleuca*).

Except for the Pacific walrus (*Odobenus rosmarus*) at least 10 specimens per species were analysed (Table S3); and for several species, exceptions of the common pattern in the presence of cusps were observed (Table 4). *MaTrics* was not designed to take intraspecific variability into account, therefore only the most common cusp patterns for each species were recorded. Deviations from the cusp patterns are present in several cusps in domestic dog, brown bear (*Ursus arctos*) and for one cusp in the red fox (*Vulpes vulpes*). Such exceptions are important as they might indicate evolutionary trends. However, variations within a species cannot be reflected in *MaTrics* as maximally one character state is given for each character representatively for a species here. Only in this way the (common) absence or presence of a trait can be compared with the genome of again one representative of a species. Studies on intraspecific variability of certain characters would need additional matrices with different intentions.

**Table 4.** Deviations in cusp patterns in the studied Carnivora. Abbreviations: M1–3, upper (indicated by number in superscript)/lower molar tooth (indicated by subscript); P^4^ – upper 4^th^ premolar

| Species | Deviation from common cusp pattern for species |
| --- | --- |
| *Canis familiaris* | Metaconid and hypoconid at M<sup>3</sup><br>Small cusp mesial of paracone at P<sup>4</sup><br>Entoconulid (mesial of entoconid) at M<sub>1</sub><br>Additional fourth lower molar |
| *Vulpes vulpes* | Small cusp mesial of paracone at P<sup>4</sup> |
| *Ursus arctos* | Second cusp palatinal at P<sup>4</sup><br>Third lingual cusp at M<sub>2</sub><br>Three metaconid-cusps at M<sub>2</sub><br>Third palatinal cusp at M<sup>1</sup> |

## Conclusion and Future Perspectives

Recent advances in molecular techniques lead to a rapid increase in the assembly and publication of genomes from various organisms. However, knowledge of the genome sequences is only a first step to understand the relationships between genomic changes, the phenotype of an organisms and phenotypic differences between different organisms (Hardison 2003). The systematic description of phenotypic information in matrix form like in *MaTrics* is necessary to understand the genome information and to deal with questions related to evolutionary biology and biomedicine. This is not restricted to mammals as the coding principles of *MaTrics*, which comply with the requirements of molecular research, can serve as a template for matrices comprising trait knowledge of other vertebrate and non-vertebrate groups. The establishment of trait matrices for various taxa could lead to a broad documentation of phenotypes for applications in comparative genomics, and, hence, enable a systematic exploration of genotype-phenotype associations.

However, trait collections such as *MaTrics* also revealed a tremendous research gap on phenotypic data. In fact, filling *MaTrics* with information on different phenotypic traits across mammals showed that detailed information on structural, physiological, or life history traits was often not available for many species, even with intensive literature research. For example, reductions of the vomeronasal system (VNS) are clearly documented in several mammals and our previous genomic comparison of 115 mammalian genomes uncovered several genes whose loss is associated with a reduced or non-functional VNS (Hecker et al. 2019a). This genomic screen also revealed that seals (Phocidae) and otters (Lutrinae) have lost some of these genes, indicating a reduced VNS. However, to the best of our knowledge, information concerning the vomeronasal organ of Phocidae and Lutrinae is not available. Indeed, the recording status in *MaTrics* for the character “vomeronasal organ” with the states absent/present is only 37%. Another example of a character, that would be assumed to be well-known, is the absence/presence of the gall bladder (“*Vesica bilaris*”), with a recording status of 70%. In other words, the recording status of the characters in the *MaTrics* demonstrate the lack of information on phenotypic traits in several species. These research gaps can only be filled by specimen-based research (e.g. Thier and Stefen 2020). Although individual studies are valuable scientific contributions, they may not suffice to close the substantial research gaps in short time. The authors see the need for more basic zoological research complementing the systematic exploration of the genomic basis of biodiversity, i.e. research activities on biodiversity genomics could be assisted by research initiatives on biodiversity phenomics (= systematically phenotyping animals in matrices like *MaTrics*).

Most of the genomic studies mentioned above identified protein coding genes associated with complex body plan changes (e.g., aquatic and aerial lifestyle of cetaceans and bats, respectively). However, evolutionary theory predicts that changes in cis-regulatory genetic elements are probably more important for morphological changes than protein-coding genes. For instance, Roscito et al. (2018) stated that the loss of morphological traits is (often) associated with the decay of the cis-regulatory elements. Consequently, the Forward Genomic approach has been further developed to include methodologies that can be successfully associate phenotypes with the loss or presence of regulatory elements (e.g., Langer et al. 2018; Langer and Hiller 2019). In awareness of these developments, the phenotype matrix presented here already provides a whole bunch of morphological characters that will be subject to further exploration in the near future. Thus, the phenotypic information compiled in *MaTrics* will be of increasing importance. This applies for instance to those referring to tooth morphology and tooth cusps discussed above. In fact, tooth characters are known to be the result of a complex signalling network involving timely graded activation and deactivation of genes controlled by regulatory elements (e.g., Jernvall and Thesleff 2000; Thesleff et al. 2001).

A last aspect to be mentioned refers to the way how phenotypic information is documented. So far, filling *MaTrics* with information is still mostly conducted by hand; experienced scientists have to control the content and to check for homology. However, some recent developments may open the door to the partial automation of this work. First, the implementation of ontologies and semantic phenotypes in the platform Morph·D·Base. The development of a respective semantic description module is already initiated (Vogt and Baum 2019; Vogt 2019). This is expected to allow the development of computer algorithms to mine data on homologous structures to establish matrices more automatically (Vogt 2018).

*MaTrics* is a new and unique data collection of phenotypic traits of mammalian species. By including homologous phenotypic traits across (an increasing number of) species, *MaTrics* and similar matrices can serve as basis for a variety of research fields as illustrated herein. The recorded phenotypic traits are well defined and fully referenced (*characters* as well the *character state* for each species). Not only literature data are accepted for the latter, but also references to specimens in collections, which contributes in a specific way to the digitalization of collection material. *MaTrics* data are directly useful in genomic studies since the *character states* are numerically coded and hence can be extracted as NEXUS file to be machine-actionable. The scientific potential of digitized phenotype matrices is apparent and motivates thinking about future development.

## Acknowledgement

We want to thank members of the German Society of Mammalogy (DGS) for stimulating discussions at the annual DGS meeting 2019 which were useful to shape the manuscript. Also, the helpful comments of the reviewers (….) are thankfully acknowledged.

## Glossary

Anatomy - “The demonstrable facts of animal structure, or also, by transference to the object, the structure or even the tissue of the animal itself.” (Snodgras 1951:173). In other words, anatomy is the part of the phenotype of an organism that refers to its physical and structural properties. At the same time, it refers to the science of anatomy, with anatomical data being facts about the anatomy of organisms.

Character Coding – The parameterized description of a quality or relation of an operational taxonomic unit.

Character Statement – see Sereno 2007

Data repository – A large database infrastructure that collects, manages, and stores data sets for data analysis, sharing and reporting. A data repository is also known as a data library or data archive. *<u>NCBI GenBank</u> is an example of a data repository for a sequence database.*

Machine actionable – Data and metadata that are structured in a formalized and consistent way so that machines (i.e. computers) can read and use them with algorithms that were programmed against this structure. Machine-actionability of data and metadata includes for instance the use of persistent identifiers for data creators (e.g. ORCIDs), organizations and funding agencies, but also open accessibility of data for machines through a corresponding application programming interface (API), and basic semantics that allow algorithms to distinguish different categories of information and apply rules to them. Machine-actionability in this sense goes beyond machine-readability which only requires data and metadata to be readable by a machine, i.e. data and metadata must be provided in a machine-readable format. Machine-readability does not necessarily require data and metadata to provide basic semantics for allowing algorithms to distinguish different categories of information contained in them.

Morphology – “Our philosophy or science of animal form, a mental concept derived from evidence based on anatomy and embryogeny, usually incapable of proof, attempting to discover structural homologies and to explain how animal organization has come to be as it is.” (Snodgrass (1951:173). In other words, morphology refers to the interpretations of anatomical facts within theories and hypotheses such as homology.

NEXUS file – A file format widely used in bioinformatics. It stores information about taxa, phenotypic characters, trees, and other information relevant for phylogenetics. Several phylogenetic programs such as PAUP, MrBayes, and Mac Clade use this format.

Phenotypic trait – A particular part of the phenotype of an organism. The Phenotype of an organism refers to its observable constituents, properties, and relations that can be considered to result from the interaction of the organism’s genotype with itself and its environment. Anatomy is the part of the phenotype that refers to the physical and structural properties of the organism.

Ontology – Ontologies are dictionaries that can be used for describing a certain reality. They consist of labeled classes and relations between classes, both with clear definitions that are ideally created by experts through consensus and that are formulated in a highly formalized canonical syntax and standardized format with the goal to yield a lexical or taxonomic framework for knowledge representation (Smith 2003). Each ontology class and relation (also called property) possesses its own Uniform Resource Identifier (URI*) through which it can be identified and individually referenced. Ontologies contain expert-curated domain knowledge about specific kinds of entities together with their properties and relations in the form of classes defined through universal statements (Schulz et al. 2009, Schulz and Jansen 2013). Ontologies in this sense do not include statements about particular entities (i.e., empirical data). (Vogt et al. 2019)

URI – A Uniform Resource Identifier (URI) is a string of characters that follows a specific structure and unambiguously identifies a particular resource. The URI can also serve as a URL (web address), and can be resolved to an IP address (see the example URI below). http://purl.obolibrary.org/obo/CL_0000255 (*for eukaryotic cell*)

## Supplementary Material

**Table S1.**
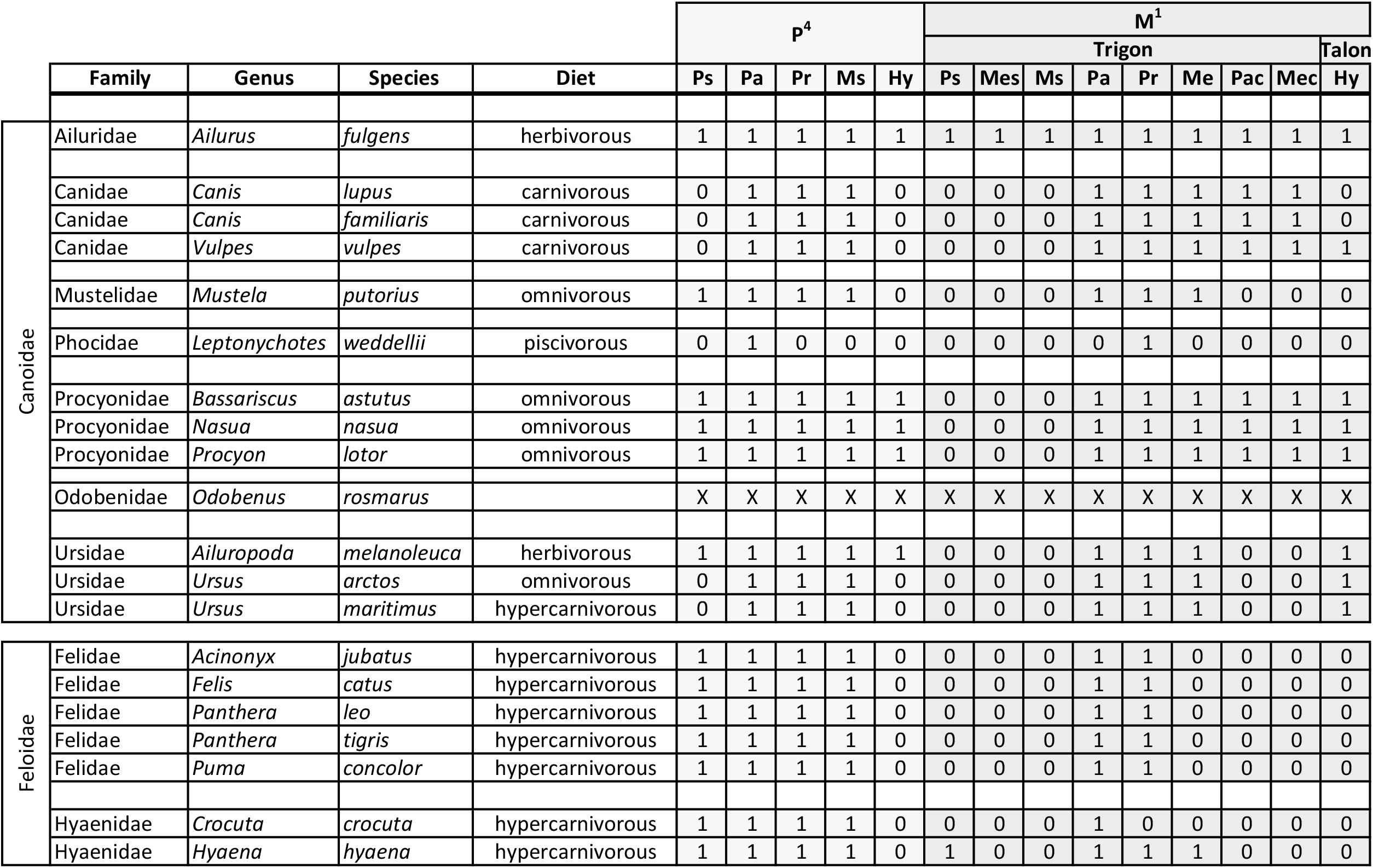
List of the mammal species allocated to order, sorted alphabetically, included in *MaTrics* so far (as of December 2020)

**Table S2.** Recording status of *MaTrics* as of December 2020. A) recording progress of the 30,282 cells for the specific character traits (absent/present, a/p; or multistate, m) as well as *missing* and *inapplicable*. Missing is the default setting and can mean a) the cell has not been treated, the information on the character state for the taxon is not known, or the information is known, but currently not retrievable (for example a specimen is known in a distant collection). The number of relevant cells as well as the percentage is given. B) Recording progress of the 206 characters for the 147 mammalian species included in *MaTrics*. The table lists the number of cells which are recorded to 100% (i.e., for all species), and to at least 75% and 50% of the species, respectively.

| *Character states* | n | % |
| --- | --- | --- |
| Recorded data (a/p and m) | 18,389 | 60.7 |
| *missing* | 8,982 | 29.7 |
| *inapplicable* | 2,911 | 9.6 |

**B**
| Recording progress (147 species) | n (traits) | % (of total number of traits) |
| --- | --- | --- |
| 100% (all species) | 1 | 0.5 |
| ≥ 75% (≥ 111 species) | 71 | 34 |
| ≥ 50% (≥ 74 species) | 129 | 63 |
| < 50% (<74 species) | 77 | 37 |

Supplementary Material document S3 Brief description of statistical methods, samples and observed *p*-values mentioned in the text

Supplementary Material Table S4 Species and the assigned material studied in the different collections (SNSD – Senckenberg Naturhistorische Sammlungen Dresden, MfN – Museum für Naturkunde Berlin) for the test study on Carnivora.

Supplementary Material document S5 Description of the tooth cusp patterns in 20 selected Carnivora.

Supplementary Material Table S6 Absence (0) and presence (1) of the analyzed cusps in the studied teeth of the carnivoran species

