## Supplementary Table S1 for "Phenotyping in the era of genomics: *MaTrics* – a digital character matrix to document mammalian phenotypic traits coded numerically"

Supplementary Material Table S1

List of the mammal species allocated to subclass, infraclass and order following Wilson and Reeder (2005) and Vaugahn (2015), sorted alphabetically, included in the *MaTrics* so far (as of June 4^th^ 2020). Breeds of domestic species are given in parentheses.

| **Number of taxon in *MaTrics*** | **Subclass: Infraclass and Order** | ***Species*** |
| --- | --- | --- |
|  | **Prototheria** |  |
| 01 | Monotremata | *Ornithorhynchus anatinus* |
| 02 | Monotremata | *Tachyglossus aculeatus* |
|  | **Theria: Metatheria** |  |
| 04 | Microbiotheria | *Dromiciops gliroides* |
| 05 | Diprotodontia | *Macropus eugenii* |
| 03 | Didelphimorphia | *Monodelphis domestica* |
| 137 | Diprotodontia | *Phascolarctos cinereus* |
| 06 | Dasyuromorphia | *Sarcophilus harrisii* |
|  | **Eutheria** |  |
| 10 | **Afrosoricida** | *Chrysochloris asiatica* |
| 09 | Afrosoricida | *Echinops telfairi* |
| 36 | **Artiodactyla** | *Bison bison* |
| 35 | Artiodactyla | *Bos grunniens mutus* |
| 33 | Artiodactyla | *Bos taurus* (breed: Hereford) |
| 34 | Artiodactyla | *Bos taurus indicus* |
| 37 | Artiodactyla | *Bubalus bubalis* |
| 46 | Artiodactyla | *Camelus bactrianus* |
| 44 | Artiodactyla | *Camelus dromedarius* |
| 117 | Artiodactyla | *Camelus ferus* (synonym to *C. batrianus* Wilson and Reeder 2005); sequenced is the feral form |
| 40 | Artiodactyla | *Capra hircus* (breed: San Clemente) |
| 119 | Artiodactyla | *Cervus elaphus* |
| 41 | Artiodactyla | *Connochaetes taurinus* |
| 48 | Artiodactyla | *Elaphurus davidianus* |
| 47 | Artiodactyla | *Giraffa camelopardalis* |
| 51 | Artiodactyla | *Muntiacus muntjak* |
| 50 | Artiodactyla | *Muntiacus reevesi* |
| 49 | Artiodactyla | *Odocoileus virginianus* |
| 39 | Artiodactyla | *Oryx gazella* |
| 38 | Artiodactyla | *Ovis aries* (breed: Texel) |
| 135 | Artiodactyla | *Ovis canadensis* |
| 42 | Artiodactyla | *Pantholops hodgsonii* |
| 43 | Artiodactyla | *Sus scrofa* (breed: Duroc) |
| 52 | Artiodactyla | *Tragulus napu* |
| 45 | Artiodactyla | *Vicugna pacos* |
| 26 | **Carnivora** | *Acinonyx jubatus* |
| 21 | Carnivora | *Ailuropoda melanoleuca* |
| 113 | Carnivora | *Ailurus fulgens* |
| 19 | Carnivora | *Canis lupus familiaris* (breed: Boxer) |
| 28 | Carnivora | *Crocuta crocuta* |
| 123 | Carnivora | *Enhydra lutris* |
| 23 | Carnivora | *Felis catus* (breed: Abyssinian) |
| 30 | Carnivora | *Leptonychotes weddellii* |
| 126 | Carnivora | *Lycaon pictus* |
| 29 | Carnivora | *Mustela putorius furo* |
| 145 | Carnivora | *Neomonachus schauinslandi* |
| 31 | Carnivora | *Odobenus rosmarus* |
| 25 | Carnivora | *Panthera leo* |
| 136 | Carnivora | *Panthera pardus* |
| 24 | Carnivora | *Panthera tigris* |
| 27 | Carnivora | *Puma concolor* |
| 22 | Carnivora | *Ursus maritimus* |
| 20 | Carnivora | *Vulpes vulpes* |
| 115 | **Cetacea** | *Balaena mysticetus* |
| 58 | Cetacea | *Balaenoptera acutorostrata* |
| 122 | Cetacea | *Delphinapterus leucas* |
| 56 | Cetacea | *Lipotes vexillifer* |
| 53 | Cetacea | *Orcinus orca* |
| 57 | Cetacea | *Physeter macrocephalus* |
| 55 | Cetacea | *Sousa chinensis* |
| 54 | Cetacea | *Tursiops truncatus* |
| 68 | **Chiroptera** | *Eidolon helvum* |
| 65 | Chiroptera | *Eptesicus fuscus* |
| 125 | Chiroptera | *Hipposideros armiger* |
| 69 | Chiroptera | *Megaderma lyra* |
| 139 | Chiroptera | *Miniopterus natalensis* |
| 64 | Chiroptera | *Myotis brandti* |
| 63 | Chiroptera | *Myotis davidii* |
| 62 | Chiroptera | *Myotis lucifugus* |
| 70 | Chiroptera | *Pteronotus parnellii* |
| 67 | Chiroptera | *Pteropus alecto* |
| 66 | Chiroptera | *Pteropus vampyrus* |
| 71 | Chiroptera | *Rhinolophus ferrumequinum* |
| 142 | Chiroptera | *Rhinolophus sinicus* |
| 144 | Chiroptera | *Rousettus aegyptiacus* |
| 72 | **Dermoptera** | *Desmodus rotundus* |
| 73 | Dermoptera | *Galeopterus variegatus* |
| 16 | **Erinaceomorpha** | *Erinaceus europaeus* |
| 13 | **Hyracoidea** | *Procavia capensis* |
| 111 | **Lagomorpha** | *Ochotona princeps* |
| 112 | Lagomorpha | *Oryctolagus cuniculus* |
| 11 | **Macroscelidea** | *Elephantulus edwardii* |
| 59 | **Perissodactyla** | *Ceratotherium simum* |
| 147 | Perissodactyla | *Dicerorhinus sumatrensis* |
| 124 | Perissodactyla | *Equus asinus* |
| 60 | Perissodactyla | *Equus caballus* |
| 61 | Perissodactyla | *Equus przewalskii* |
| 129 | **Pholidota** | *Manis javanica* |
| 32 | Pholidota | *Manis pentadactyla* |
| 114 | **Primates** | *Aotus nancymaae* |
| 81 | Primates | *Callithrix jacchus* |
| 76 | Primates | *Carlito syrichta syrichta* |
| 118 | Primates | *Cebus capucinus* |
| 138 | Primates | *Cercocebus atys* |
| 92 | Primates | *Chlorocebus sabaeus* |
| 120 | Primates | *Colobus angolensis* |
| 77 | Primates | *Daubentonia madagascariensis* |
| 85 | Primates | *Gorilla gorilla* |
| 82 | Primates | *Homo sapiens* |
| 89 | Primates | *Macaca fascicularis* |
| 88 | Primates | *Macaca mulatta* |
| 127 | Primates | *Macaca nemestrina* |
| 90 | Primates | *Macaca thibetana* |
| 128 | Primates | *Mandrillus leucophaeus* |
| 79 | Primates | *Microcebus murinus* |
| 134 | Primates | *Nasalis larvatus* |
| 87 | Primates | *Nomascus leucogenys* |
| 78 | Primates | *Otolemur garnettii* |
| 86 | Primates | *Pan paniscus* |
| 83 | Primates | *Pan troglodytes* |
| 93 | Primates | *Papio anubis* |
| 140 | Primates | *Piliocolobus tephrosceles* |
| 84 | Primates | *Pongo abelii* |
| 141 | Primates | *Propithecus coquereli* |
| 143 | Primates | *Rhinopithecus bieti* |
| 91 | Primates | *Rhinopithecus roxellana* |
| 80 | Primates | *Saimiri boliviensis* |
| 146 | **Proboscidea** | *Elephas maximus* |
| 14 | Proboscidea | *Loxodonta africana* |
| 116 | **Rodentia** | *Castor canadensis* |
| 97 | Rodentia | *Cavia aperea* |
| 96 | Rodentia | *Cavia porcellus* |
| 108 | Rodentia | *Chinchilla lanigera* |
| 98 | Rodentia | *Cricetulus barabensis* |
| 121 | Rodentia | *Cricetulus griseus* |
| 104 | Rodentia | *Cryptomys damarensis* |
| 102 | Rodentia | *Dipodomys ordii* |
| 103 | Rodentia | *Heterocephalus glaber* |
| 105 | Rodentia | *Ictidomys tridecemlineatus* |
| 109 | Rodentia | *Jaculus jaculus* |
| 130 | Rodentia | *Marmota marmota* |
| 131 | Rodentia | *Meriones unguiculatus* |
| 99 | Rodentia | *Mesocricetus auratus* |
| 100 | Rodentia | *Microtus ochrogaster* |
| 132 | Rodentia | *Mus caroli* |
| 94 | Rodentia | *Mus musculus* |
| 133 | Rodentia | *Mus pahari* |
| 110 | Rodentia | *Octodon degus* |
| 101 | Rodentia | *Peromyscus maniculatus* |
| 95 | Rodentia | *Rattus norvegicus* |
| 107 | Rodentia | *Spalax galili* |
| 106 | Rodentia | *Spermophilus dauricus* |
| 74 | **Scandentia** | *Tupaia belangeri* |
| 75 | Scandentia | *Tupaia belangeri chinensis* |
| 15 | **Sirenia** | *Trichechus manatus* |
| 18 | **Soricomorpha** | *Condylura cristata* |
| 17 | Soricomorpha | *Sorex araneus* |
| 12 | **Tubulidentata** | *Orycteropus afer* |
| 08 | **Xenarthra** | *Choloepus hoffmanni* |
| 07 | Xenarthra | *Dasypus novemcinctus* |
