## Supplementary Document S3 for "Phenotyping in the era of genomics: *MaTrics* – a digital character matrix to document mammalian phenotypic traits coded numerically"

**Supplementary Material Document S3** Brief description of statistical methods, samples and observed *p*-values mentioned in the text

Peggy Wehner, Matthias Rudolf

Seventeen morphological characters of the gastrointestinal tract and dentition given in *MaTrics* were tested in order to elucidate significant differences between a) the two dietary groups herbivores and carnivores as defined in the list of Hecker et al. (2019; see below) and b) the absence or presence of PNLIPRP1 and NR1I3, respectively. Additionally, it was tested whether a significant relation to one of the mentioned genes is present within carnivorous or within herbivorous species. Fisher’s exact test was used for nominal scaled variables and exact biserial rank correlations for ordinal scaled variables, both on the 5% probability level. For the analyses the program R was used. The results are listed in the tables below. It is not within the scope of this paper to go into details here, neither extending statistical tests nor interpreting the given results in detail (as they are part of an on-going project). Nevertheless, the statistical methods and results presented here highlight the potential of their use with *MaTrics*.

The *p*-values are given below for the tests of 17 characters. The scale level of the characters is given in brackets. The reported *p*-values in bold indicate statistical significance. No correction was made for multiple testing, so the risk of accumulation of alpha errors should be considered when interpreting the results represented.

| character | *p*-value in relation to diet category | *p*-value in relation to PNLIPRP1 | *p*-value in relation to NR1I3 |
| --- | --- | --- | --- |
| Vesica bilaris (nominal) | 0.7411 | 0.4951 | **0.0028** |
| Caecum (nominal) | **0.0003** | **0.0051** | **0.0056** |
| Appendix vermiformis (nominal) | 0.5929 | 0.4920 | 0.5576 |
| Diet with heightened fat content (nominal) | **0.0000** | **0.0000** | **0.0193** |
| Xenobiotics (nominal) | **0.0000** | **0.0002** | **0.0000** |
| Canine (sup) (nominal) | **0.0002** | **0.0000** | **0.0377** |
| Canine (inf) (nominal) | 0.1107 | 0.1809 | 0.3964 |
| Cheek teeth (sup) / number (ordinal) | 0.6142 | 0.5293 | 0.5381 |
| Cheek teeth (inf) / number (ordinal) | 0.2492 | 0.2561 | 0.4107 |
| Homodont (ordinal) | 0.1037 | 0.8204 | **0.0075** |
| Canine / relative height in relation to occlusal level of tooth row (sup) (ordinal) | 0.1626 | 1.0000 | 0.6571 |
| Canine / relative height in relation to occlusal level of tooth row (inf) (ordinal) | 0.3883 | 0.4667 | 0.0696 |
| Number functional incisors (sup) (ordinal) | **0.0159** | **0.0000** | 0.3976 |
| Number functional incisors (inf) (ordinal) | 0.7162 | 0.1348 | 0.3531 |
| Premolares (sup/inf), brachydont (nominal) | **0.0001** | 0.0649 | 0.0571 |
| Molares (sup/inf), brachydont (nominal) | **0.0000** | **0.0086** | **0.0209** |
| Molares (sup/inf) / occlusal pattern (nominal) | **0.0000** | **0.0000** | **0.0000** |

| character | *p*-value in relation to PNLIPRP1 |  | *p*-value in relation to dietary groups |  |
| --- | --- | --- | --- | --- |
|  | herbivores | carnivores | PNLIPRP1 absent | PNLIPRP1 present |
| Vesica bilaris | 1.0000 | **0.0361** | **0.0429** | 1.0000 |
| Caecum | 1.0000 | 0.5282 | 0.2982 | 0.0686 |
| Appendix vermiformis | 0.1333 | 1.0000 |  | 0.2047 |
| Diet |  | 0.2092 | **0.0074** | **0.0008** |
| Diet with heightened fat content | 0.0593 | 0.1200 | 0.1553 | 0.1538 |
| Canine (sup) | 0.1793 | 0.1765 | 1.0000 | 0.1620 |
| Canine (inf) | 1.0000 | 0.1765 | 0.4286 | 0.1620 |
| Cheek teeth (sup) / number | 1.0000 |  |  | 1.0000 |
| Cheek teeth (inf) / number | 1.0000 |  |  | 0.9729 |
| Homodont | 1.0000 | 0.2381 | 0.1333 | 0.4097 |
| Canine / relative height in relation to occlusal level of tooth row (sup) | 0.3333 |  |  | 0.0659 |
| Canine / relative height in relation to occlusal level of tooth row (inf) | 0.3333 |  |  | 0.0513 |
| Number functional incisors (sup) | **0.0073** | **0.0351** | 0.4048 | 0.9098 |
| Number functional incisors (inf) | 0.6379 | 0.0585 | **0.0476** | 0.9609 |
| Premolares (sup/inf), brachydont | 0.4909 |  |  | **0.0018** |
| Molares (sup/inf), brachydont | 0.5412 |  |  | **0.0018** |
| Molares (sup/inf) / occlusal pattern | 1.0000 |  |  | **0.0004** |

Hecker et al. (2019) used 52 placental mammals covering 11 orders of 3 superorders listed here assigned taxonomically: Glires: **Rodentia** (Bathyergidae: *Heterocephalus glaber*, *Cryptomys damarensis*; Caviomorpha: Caviidae: *Cavia porcellus*, *Cavia aperea*; Octodontidae: *Octodon degus*) and **Lagomorpha** (Leporidae: *Oryctolagus cuniculus*; Octodontidae: *Ochotona princeps*); Laurasiatheria: **Artiodactyla** (terrestrial Artiodactyla: *Vicugna pacos*, Bovidae: *Bos taurus*, *Bos taurus* *indicus*, *Bison bison*, *Bos grunniens* *mutus*, *Bubalus bubalis*, *Ovis aries*, *Ovis canadensis*, *Capra hircus*, *Pantholops hodgsonii*); **Cetacea**: Odontoceti: *Tursiops* *truncatus*, *Orcinus orca*, *Delphinapterus leucas*, *Lipotes vexillifer*, *Physeter macrocephalus*; Mysticeti: *Balaenoptera acutorostrata*, *Balaena mysticetus*); **Perissodactyla** (Equidae: *Equus caballus*, *Equus asinus*; Rhinocerotidae: *Ceratotherium* *simum*, *Dicerorhinus sumatrensis*); **Carnivora** (*Felis catus*, *Acinonyx jubatus*, *Panthera* *tigris*, *Panthera pardus*, *Mustela putorius furo*, *Enhydra lutris*, *Odobenus rosmarus*, *Leptionychotes weddellii*, *Neomonachus schauinslandi*) and **Pholidota** (*Manis* *pentadactyla*, *Manis javanica*) (both orders united to Ferae); **Chiroptera** (Pteropodidae: *Pteropus alecto*, *Pteropys vampyrus*, *Rousettus aegyptiacus*; Vespertilionidae: *Eptesicus fuscus*, *Myotis davidii*, *Myotis brandti*, *Myotis lucifugus*); **Lipotyphla** (*Sorex araneus*, *Condylura cristata*); Afrotheria: **Proboscidea** (Elephantidae: *Loxodonta* *africana*, *Elephas maximus*) and **Sirenia** (*Trichechus manatus*) (both orders united to Paenungulata); **Afrosoricida** (*Chrysochloris asiatica*).
