## Supplementary Table S4 for "Phenotyping in the era of genomics: *MaTrics* – a digital character matrix to document mammalian phenotypic traits coded numerically"

Table S4 Species and the assigned material studied in the different collections (SNSD – Senckenberg Naturhistorische Sammlungen Dresden, MfN – Museum für Naturkunde Berlin) for the test study on Carnivora.

| Ailuridae | *Ailurus fulgens* | SNSD | B9889, B9903, B10124, B17173, B17477, B17478, B19080 |
| --- | --- | --- | --- |
| Ailuridae | *Ailurus fulgens* | MfN | ZMB_Mam_3256, ZMB_Mam_13346, ZMB_Mam_45250 |
| Canidae | *Canis lupus* | SNSD | B1174, B2473, B3341, B4956, B4957, B5721, B8303, B8304, B19334, B22149 |
|  | *Canis lupus familiaris* | SNSD | B4558, B4679, B4680, B5063, B8861, B8862, B9573, B14230, B14234, B14236, B14238, B14241, B14242, B14268, B14269, B14287, B14293, B14508, B14509, B14781, B16276, B16277, B16281, B16285, B16287, B16291, B16292, B16295, B16296, B16298, B16300, B16301, B16303, B16304, B16305, B16307, B16308, B18643, B18653, B18654, B19399, B22161, B22162, B22163, B22164, B22168, B22480B, B22481B, B22485, B22577 |
|  | *Vulpes vulpes* | SNSD | B5341, B5789, B6538, B6542, B7082, B7348, B7350, B7379, B7969, B7972, B8329, B8335, B8827, B8849, B8850, B8851, B8867, B8905, B9500, B9566, B9567, B9780, B9781, B10351, B10352, B12756, B13047, B13048, B14875, B15057, B15058, B15060, B15064, B16273, B16274, B18646, B18647, B19070, B19456, B19495, B22477, B22478A, B22478B, B22478C, B22588, B22589, B24694, B26172, B27007, B27694 |
| Felidae | *Acinonyx jubatus* | SNSD Mfn | B5415 (1), B5415 (2), B6189, B16434 ZMB_Mam_12725, ZMB_Mam_16812, ZMB_Mam_56285, ZMB_Mam_56308, ZMB_Mam_82181, ZMB_Mam_82977 |
|  | *Felis catus* | SNSD | B4230, B8858, B8860, B9318, B15641, B22288, B22289, B22292, B22294, B22474 |
|  | *Panthera leo* | SNSD | B1504, B4366, B5578, B6281, B6632, B9816, B9817, B12826, B12827, B12828 |
|  | *Panthera tigris* | SNSD | B2169, B2170, B7470, B16438, B16441, B16443, B16561, B16733, B16747, B22863 |
|  | *Puma concolor* | SNSD | 936, B2435, B5720, B6166, B10021, B16455, B16457, B16459, B17467, B25769 |
| Hyaenidae | *Crocuta crocuta* | SNSD | B4936, B5096, B8154, B8155, B8156, B9819, B14171, B16003, B16444, B16445 |
|  | *Hyaena hyaena* | SNSD |  |
|  | *Hyaena hyaena* | MfN | ZMB_Mam_14824, ZMB_Mam_14826, ZMB_Mam_31303, ZMB_Mam_82301, ZMB_Mam_82336, ZMB_Mam_82337, ZMB_Mam_82345 |
| Mustelidae | *Mustela putorius* | SNSD | B191/246, B1265, B4647, B4901, B5615, B8730, B8870, B8873, B9473, B9562 |
| Odobenidae | *Odobenus rosmarus* | SNSD | none, B3235, B3357, B22137 |
| Phocidae | *Leptonychotes weddelli* | SNSD | B5029 |
| Phocidae | *Leptonychotes*  *weddellii* | MfN | ZMB_Mam_20275, ZMB_Mam_27674, ZMB_Mam_36277, ZMB_Mam_36278, ZMB_Mam_36285, ZMB_Mam_36290, ZMB_Mam_36294, ZMB_Mam_86627, ZMB_Mam_86628 |
| Procyonidae | *Bassariscus astatus* | SNSD | B7086, B16473 |
|  | *Bassariscus astatus* | SNSD | B1855, B2420, B4655, B6174, B6313, B8832, B11873, B16488, B16498, B22502, B22594, B22618 |
|  | *Nasua nasua* | SNSD | B1855, B2420, B4655, B6174, B6313, B8832, B11873, B16488, B16498, B22502, B22594, B22618 |
|  | *Procyon lotor* | SNSD | B2399, B6701, B22583, B27695 |
|  | *Procyon lotor* | MfN | ZMB_Mam_571, ZMB_Mam_2849, ZMB_Mam_10218, ZMB_Mam_56630, ZMB, Mam_77056, ZMB_Mam_77156 |
| Ursidae | *Ailuropoda melanoleuca* | MfN | ZMB_Mam_17246, ZMB_Mam_17542, ZMB_Mam_37026, ZMB_Mam_85761 |
|  | *Ursus arctos* | SNSD | A1271, B1864, B2194, B3353, B3355, B5234, B8308, B8309, B13257, B16334, B16335, B16336, B16337, B16338, B16339, B16369, B16370, B16371, B16372, B16373, B21795, B22620, B22621, B22622, B22623, B24691, B27909 |
|  | *Ursus maritimus* | SNSD | B2428, B4550, B8419, B13255, B13256, B15669, B15670, B16374, B16375, |
