## Supplementary Document S5 for "Phenotyping in the era of genomics: *MaTrics* – a digital character matrix to document mammalian phenotypic traits coded numerically"

Description to tooth cusp patterns in 20 selected Carnivora

Content:

Ailuridae	*Ailurus fulgen*s

Canidae	*Canis lupus*

*Canis lupus familiaris*

*Vulpes vulpes*

Felidae	*Acinonyx jubatus*

*Felis catus*

*Panthera leo*

*Panthera tigris*

*Puma concolor*

Hyaenidae	*Crocuta crocuta*

*Hyaena hyaena*

Mustelidae	*Mustela putorius*

Procyonidae	*Bassariscus astatus*

*Nasua nasua*

*Procyon lotor*

Ursidae	*Ailuropoda melanoleuca*

*Ursus arctos*

*Ursus maritimus*

The studied material is listed in Supplementary Material Table S4

The results are simplified in Supplementary Material Table S6

The two species, *Odobenus rosmarus (*Odobenidae) and *Leptonychotes weddellii* (Phocidae) were left out, because cusps can’t be described.

**Ailuridae**

*Ailurus fulgens* as single member of the Ailuridae is characterized by the dental formula 3132/3142.

***Ailurus fulgens* CUVIER, 1825 – red panda, lesser panda**

The rounded triangular P4 is formed by parastyle (mesiobuccal), paracone (buccal), metastyle (distobuccal), protocone (mesiolingual), and hypocone (distolingual). The M1 is triangular with rounded corners and divided into trigon and talon. The trigon is made up of the main cusps paracone (mesiobuccal), metacone (distobuccal), and protocone (mesiolingual), as well as the smaller cuspules paraconule (mesial) and metaconule (distal). At the buccal side there are the stylar cusps parastyle, mesostyle, and metastyle. The talon is formed by the hypocone (distolingual). The M2 is similar to the M1. The trigon comprises paracone (mesiobuccal), metacone (distobuccal), protocone (mesiolingual), paraconule (mesial), and metaconule (distal). At the buccal side there are the stylar cusps parastyle, mesostyle, and metastyle. A hypocone is not present. The rectangular m1 is divided into trigonid and talonid. The trigonid contains paraconid (mesial), metaconid (mesiolingual), and protoconid (mesiobuccal), while the talonid is made up of hypoconid (distobuccal), hypoconulid (distal), entoconid (distolingual), and entoconulid (lingual). The m2 is similar to the m1. Protoconid (mesiobuccal) and metaconid (mesiolingual) are forming the trigonid. The talonid contains hypoconid (distobuccal), hypoconulid (distal), entoconid (distolingual), and entoconulid (lingual).

**Canidae**

The investigated members of the family Canidae are characterized by the dental formula 3142/3143.

***Canis lupus* LINNAEUS, 1758 – wolf**

The mesiodistally elongated P4 is characterized by a high paracone-metastyle blade and the reduced protocone (mesiolingual). The triangular M1 contains the main cusps paracone (mesiobuccal), metacone (distobuccal), and protocone (mesiolingual), as well as the smaller cuspules paraconule (mesial) and metaconule (distal). At the lingual side there is a high cingulum present. The M2 is similar to the M1 but smaller. It consists of paracone (mesiobuccal), metacone (distobuccal), protocone (mesiolingual), paraconule (mesial), and metaconule (distal). A high cingulum is present at the distolingual side. The m1 is rectangular and divided into trigonid and talonid. The trigonid is characterized by a high paraconid-protoconid blade and the smaller metaconid (lingual). The talonid is made up of hypoconid (distobuccal) and entoconid (distolingual). The small and rounded m2 is also divided into trigonid and talonid. Protoconid (mesiobuccal) and metaconid (mesiolingual) are forming the trigonid, while the talonid is formed by the hypoconid (distobuccal). The rounded m3 is extremely reduced and comprises only one cuspule which could correspond to the protoconid.

***Canis lupus familiaris* (LINNAEUS, 1758) – domestic dog**

The P4 is mesiodistally elongated and contains a high paracone-metastyle blade, as well as the reduced protocone (mesiolingual). The triangular M1 is characterized by the main cusps paracone (mesiobuccal), metacone (distobuccal), and protocone (mesiolingual). Paraconule (mesial) and metaconule (distal) are smaller cuspules of the M1. The M2 is similar to the M1 but smaller. It is made up of the main cusps paracone (mesiobuccal), metacone (distobuccal), protocone (mesiolingual), and the small cuspule paraconule (mesial). The m1 is rectangular and divided into trigonid and talonid. The trigonid consists of a high paraconid-protoconid blade and the smaller metaconid (lingual). The talonid is made up of hypoconid (distobuccal) and entoconid (distolingual). The small and rounded m2 is also divided into trigonid and talonid. Protoconid (mesiobuccal) and metaconid (mesiolingual) are forming the trigonid, while the hypoconid is forming the talonid. The rounded m3 is extreme reduced and contains one cuspule which could correspond to the protoconid.

Nearly all of the analyzed specimens show the same cusp pattern. In some specimens paraconule and metaconule are extremely reduced or lost at the M1 and/or M2. A hypocone is sometimes present at the M1 but only two of 50 specimens show a hypocone at the M2. Seven of 50 specimens contain a entoconulid at the m1. In eight of 50 specimens there is an entoconid at the m2. One specimen is characterized by an additional small cusp mesial of the paracone (SNSD B14236). In one specimen an additional fourth lower molar is existing (SNSD B16291). The m3 of one specimen is made up of three cusps which could correspond to protoconid, metaconid, and hypoconid (SNSD B9573).

***Vulpes vulpes* (LINNAEUS,1758) – red fox**

The mesiodistally elongated P4 contains the high paracone-metastyle blade and the protocone (mesiolingual). The triangular M1 is divided into trigon and talon. The trigon is made up of the main cusps paracone (mesiobuccal), metacone (distobuccal), and protocone (mesiolingual), as well as the smaller cuspules paraconule (mesial) and metaconule (distal). The hypocone (distolingual) is part of the talon. The M2 is similar to the M1 but smaller. Paracone (mesiobuccal), metacone (distobuccal), protocone (mesiolingual), paraconule (mesial), and metaconule (distal) are forming the trigon, while the hypocone (distolingual) indicates the talon. The rectangular m1 is divided into trigonid and talonid. The trigonid is characterized by a high paraconid-protoconid blade and the smaller metaconid (mesiolingual). The talonid consists of hypoconid, (distobuccal), hypoconulid (distal), entoconid (distolingual), and entoconulid (lingual). The triangular m2 is also divided into trigonid and talonid. Protoconid (mesiobuccal) and metaconid (mesiolingual) are forming the trigonid, while the talonid is made up of hypoconid (distobuccal), entoconid (distolingual), and entoconulid (lingual). The rounded m3 is extreme reduced and contains two cuspules which could correspond to the protoconid (buccal) and the metaconid (lingual).

Nearly all of the analyzed specimens show the same cusp pattern. But two specimens show an additional cusp mesial of the paracone at the P4 (SNSD B8827, SNSD B10352). In some specimens the cuspules at the M2 – paraconule and metaconule – are extreme reduced or lost. In five of 50 red fox specimens there is no entoconulid at the m1 present. Five of 50 specimens show a hypoconulid at the m2. In some specimens entoconid and entoconulid of m2 are extreme reduced or lost.

**Felidae**

The investigated Felidae are characterized by elongate, blade-like carnassials and the dental formula 3131/3131.

***Acinonyx jubatus* (SCHREBER, 1775) – cheetah**

The P4 is mesiodistally elongated. Paracone (central) and metastyle (distal) are forming a high blade. Parastyle (mesiobuccal) and protocone (mesiolingual) are smaller. The M1 is reduced and contains two cuspules which could correspond to paracone (buccal) and protocone (lingual). The m1 is mesiodistally elongated and made up of the paraconid-protoconid blade. The distal cingulum could correspond to the talonid.

Nearly all of the analyzed samples show the same cusp pattern except one: at m1 of ZMB_Mam_56285 there is a third cusp distolingual to the protoconid. This could correspond to the metaconid. The reduced talonid is located at the distal end.

Paper Solé & Ladevèze, Evolution and Development, 2017, p.61

→ hypoconid as structure of the talonid, no metaconid in cats

- Gaunt: The development of the deciduous cheek teeth of the cat, 1959
- Jernvall: Mammalian molar susp pattern: Developmental mechanisms of diversity, 1995
- Tims: On the tooth-genesis in the Canidae, 1896

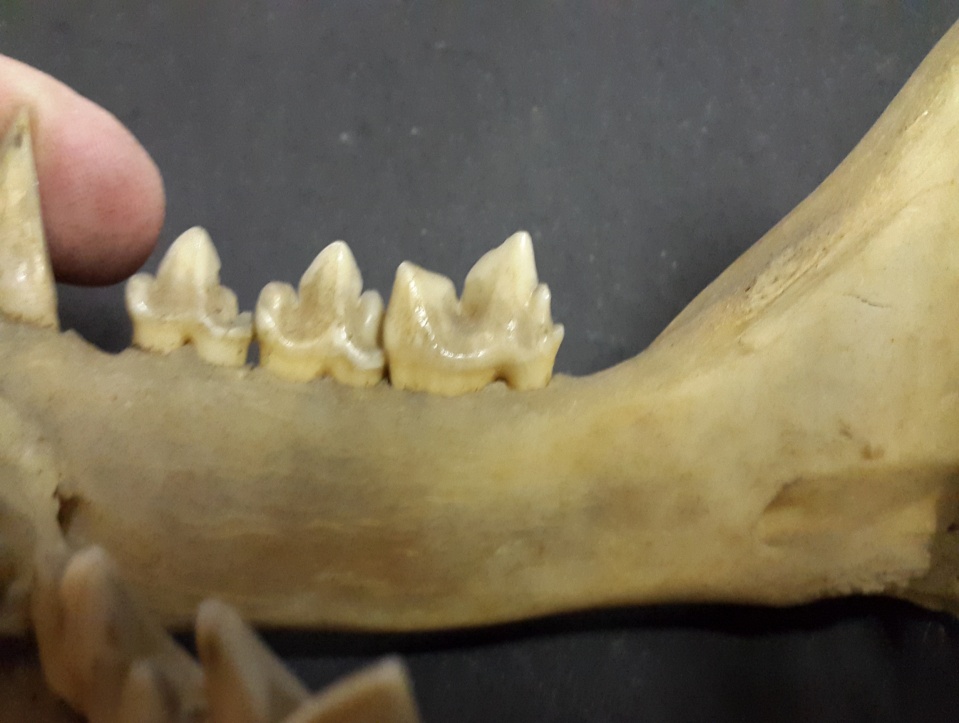

20170524_143903

med

*Acinonyx jubatus* (ZMB_Mam_56285), M1 upper jaw with metaconid

***Felis catus* LINNAEUS, 1758 – domestic cat**

The mesiodistally elongated P4 consists of the smaller cusps parastyle (mesiobuccal) and protocone (mesiolingual) and the high paracone-metastyle blade. The M1 is reduced and made up of two cuspules which could correspond to paracone (buccal) and protocone (lingual). The m1 is also mesiodistally elongated and comprises the paraconid-protoconid blade. The distal cingulum could correspond to the talonid.

***Panthera leo* (LINNAEUS, 1758) – lion**

The P4 is mesiodistally elongated. Paracone (central) and metastyle (distal) are forming a high blade while parastyle (mesiobuccal) and protocone (mesiolingual) are smaller. The M1 is reduced and contains two cuspules which could correspond to paracone (buccal) and protocone (lingual). The m1 is mesiodistally elongated and shows the paraconid-protoconid blade. There is no cingulum at the distal end.

***Panthera tigris* (LINNAEUS, 1758) – tiger**

The mesiodistally elongated P4 is characterized by a high paracone-metastyle blade. Parastyle (mesiobuccal) and protocone (mesiolingual) are smaller. The reduced M1 contains two cuspules which could correspond to paracone (buccal) and protocone (lingual). The mesiodistally elongated m1 consists of the paraconid-protoconid blade. At the distal end there is no cingulum present.

***Puma concolor* (LINNAEUS, 1771) – puma, cougar, mountain lion**

The P4 is mesiodistally elongated. It consists of paracone (central) and metastyle (distal) which are forming a high blade and the smaller cusps parastyle (mesiobuccal) and protocone (mesiolingual). The M1 is reduced. The two cuspules could correspond to the paracone (buccal) and the protocone (lingual). The m1 is also mesiodistally elongated and characterized by the paraconid-protoconid blade. There is no cingulum at the distal end.

**Hyaenidae**

Members of the Hyaenidae (except *P. cristata*) are characterized by elongate, bladelike carnassials and the dental formula 3141/3131.

|  | P4 | | | | M1 | | | | m1 | | | | | |
| --- | --- | --- | --- | --- | --- | --- | --- | --- | --- | --- | --- | --- | --- | --- |
|  | Pa | Pr | Ps | Ms | Pa | Pr | Ps | Me | pad | prd | med | talonid | hyd | end |
| *C. crocuta* | + | + | + | + | ? | ? | - | - | + | + | - | + | - | - |
| *H. hyaena* | + | + | + | + | + | + | + | + | + | + | + | + | + | + |

+ present, - absent, ? not defined

***Crocuta crocuta* (ERXLEBEN, 1777) – spotted hyena**

The P4 is mesiodistally elongated. Paracone (central) and metastyle (distal) are forming a high blade while parastyle (mesiobuccal) and protocone (mesiolingual) are smaller. The M1 is extreme reduced and moved lingually. The mesiodistally elongated m1 is characterized by the paraconid-protoconid blade. The cingulum at the distal end could correspond to the talonid.

***Hyaena hyaena* (LINNAEUS, 1758) – striped hyena**

The mesiodistally elongated P4 consists of the paracone-metastyle blade and the smaller cusps parastyle (mesiobuccal) and protocone (mesiolingual). The M1 is reduced. It is made up of the parastyle (mesiobuccal), paracone (buccal), metacone (distobuccal), and protocone (lingual). The m1 is also mesiodistally elongated and is divided into trigonid and talonid. The paraconid-protoconid blade and the metaconid (lingual) are forming the trigonid. The small talonid is made up of hypoconid (distobuccal) and entoconid (distolingual).

**Mustelidae**

The dental formula of the Mustelidae is variable. *Mustela putorius* is one example which shows the dental formula 3131/3132.

***Mustela putorius* LINNAEUS, 1758 – European polecat**

The P4 is mesiodistally elongated with a high paracone-metastyle blade. Parastyle (mesiobuccal) and protocone (mesiolingual) are smaller. The M1 is wide buccolingually and in occlusal view it has the shape of an hourglass. Paracone (mesiobuccal) and metacone (distobuccal) are situated at the buccal side, while the lingual side shows the protocone. The m1 is mesiodistally elongated and divided into trigonid and talonid. The trigonid is made up of the paraconid-protoconid blade, there is no metaconid. The talonid is reduced with only the small cuspule of the hypoconid (distobuccal). The m2 is rounded and extreme reduced. The two cuspules could correspond to paraconid (buccal) and protoconid (lingual).

**Procyonidae**

The investigated members of the Procyonidae are characterized by the dental formula 3142/3142.

***Bassariscus astutus* (LICHTENSTEIN, 1830) – ringtail**

The P4 is characterized by a high paracone-metastyle blade. At the lingual side there are the smaller cusps protocone (mesiolingual) and hypocone (distolingual). A parastyle is not present. The triangular M1 is divided into trigon and talon. Paracone (mesiobuccal), metacone (distobuccal), and protocone (mesiolingual) are the main cusps of the trigon. Two smaller cuspules - paraconule (mesial) and metaconule (distal) - are situated between paracone and protocone and metacone and protocone, respectively. The distolingual hypocone represents the talon. The M2 is smaller than the M1 but looks similar. The trigon is made up of paracone (mesiobuccal), metacone (distobuccal), protocone (mesiolingual), paraconule (mesial), and metaconule (distal). There is no hypocone existing. The m1 is divided into trigonid and talonid. The trigonid is made up of a paraconid-protoconid blade and the metaconid. The talonid consists of hypoconid (distobuccal) and entoconid (distolingual). The m2 is smaller than the m1 and also divided into trigonid and talonid. The trigonid is made up of protoconid (mesiobuccal) and metaconid (mesiolingual), while the talonid contains hypoconid (distobuccal) and entoconid (distolingual).

Nearly all of the 10 analyzed specimens show the same cusp pattern. But there are some differences: some cusps were added while others were lost. The hypocone at the P4 is only present in seven of 10 specimens. Only one individual ZMB_Mam_1634.5 has a hypocone at the M2. The m1 of six of 10 specimens comprises a hypoconulid and in four of 10 specimens an entoconulid. Only four of 10 specimens show an entoconulid at the m2.

***Nasua nasua* (LINNAEUS, 1766) – South American coati**

The P4 is triangular. At the buccal side it is made up of the parastyle (mesiobuccal), paracone (buccal), and an extremely reduced metastyle (distobuccal). The lingual side contains protocone (mesiolingual) and hypocone (distolingual). The M1 is also triangular and divided into trigon and talon. The trigon is made up of paracone (mesiobuccal), metacone (distobuccal), protocone (mesiolingual), and the smaller cuspules paraconule (mesial) and metaconule (distal). The hypocone (distolingual) contributes to the talon. The M2 looks nearly similar to the M1 and has the same size. The trigon is formed by paracone (mesiobuccal), metacone (distobuccal), protocone (lingual), and the smaller cuspules paraconule (mesial) and metaconule (distal). But there is no talon present. The rectangular m1 is divided into trigonid and talonid. The trigonid consists of paraconid (mesiolingual), protoconid (mesiobuccal), and metaconid (lingual). The talonid is built by the hypoconid (distobuccal) and entoconid (distolingual). The m2 is also divided into trigonid and talonid. Protoconid (mesiobuccal) and metaconid (mesiolingual) are forming the trigonid while hypoconid (distobuccal), hypoconulid (distal), entoconid (distolingual), and entoconulid (lingual) are forming the talonid.

Nearly all of the analyzed specimens show the same cusp pattern. But there are some differences: some cusps were added while others were lost. Three of 12 specimens show a hypoconulid at the m1, one of 12 an entoconulid. One of 12 specimens lost the hypoconulid at the m2, whereas six of 12 show an entoconulid at the m2.

***Procyon lotor* (LINNAEUS, 1758) – racoon**

The rounded P4 consists of parastyle (mesiobuccal), paracone (buccal), and an extremely reduced metastyle (distobuccal). It is completed by the protocone (mesiolingual) and hypocone (distolingual). The M1 is rounded and divided into trigon and talon. The trigon is made up of paracone (mesiobuccal), metacone (distobuccal), protocone (mesiolingual), and the smaller cuspules paraconule (mesial) and metaconule (distal). The talon is made up of the hypocone (distolingual). The triangular M2 only consists of the trigon cusps paracone (mesiobuccal), metacone (distobuccal), protocone (lingual), and the smaller cuspules paraconule (mesial) and metaconule (distal). The rectangular m1 is divided into trigonid and talonid. Paraconid (mesiolingual), metaconid (lingual), and protoconid (mesiobuccal) form the trigonid, whereas hypoconid (distobuccal), hypoconulid (distal), entoconid (distolingual), and entoconulid (lingual) form the talonid. The m2 is made up of the trigonid cusps protoconid (mesiobuccal) and metaconid (mesiolingual), as well as the talonid cusps hypoconid (distobuccal), hypoconulid (distal), entoconid (distolingual), and entoconulid (lingual).

Nearly all of the analyzed specimens show the same cusp pattern but some are different. In five of 10 specimens there is no hypoconulid at the m1, in two of 10 no entoconulid. One specimen lost the hypoconulid at the m2, two the entoconulid.

**Ursidae**

The dental formula of the investigated Ursidae is 3142/3143.

***Ailuropoda melanoleuca* (DAVID, 1869) - giant panda**

The triangular P4 consists of parastyle (mesiobuccal), paracone (buccal), and metastyle (distobuccal) as well as protocone (mesiolingual) and hypocone (distolingual). The rounded rectangular M1 contains paracone (mesiobuccal), metacone (distobuccal), and protocone (mesiolingual) as part of the trigon and hypocone (distolingual) as part of the talon. The M2 is nearly similar to the M1 except the elongated talon. The buccal side is made up of paracone (mesiobuccal) and metacone (distobuccal), while the lingual side is made up of protocone (mesiolingual) and hypocone (distolingual). Next to these main cusps there are a lot of smaller cuspules present on the surface. The rectangular m1 is divided into trigonid and talonid. The trigonid consists of paraconid (mesiolingual), metaconid (lingual), and protoconid (mesiobuccal). Hypoconid (distobuccal), hypoconulid (distal), and entoconid (distolingual) are forming the talonid. The m2 is also divided into trigonid and talonid. Paraconid (mesiolingual) and protoconid (mesiobuccal) are forming the trigonid, while hypoconid (distobuccal), hypoconulid (distal), and entoconid (distolingual) are forming the talonid. There are several smaller cuspules on the surface of m2. The m3 is triangular, extremely wrinkled and there are no defined cusps.

Nearly all of the four analyzed specimens show the same cusp pattern. Only one specimen lost the hypoconulid at m2.

***Ursus arctos* LINNAEUS, 1758 – brown bear**

The nearly triangular P4 shows a distinct paracone (mesiobuccal), metastyle (distobuccal), and protocone (lingual). The rounded rectangular M1 is divided into trigon and talon. The trigon is made up of paracone (mesiobuccal), metacone (distobuccal), and protocone (mesiolingual). The hypocone (distolingual) contributes to the talon. The M2 is nearly similar to the M1 except the elongated talon. Paracone (mesiobuccal), metacone (buccal), and protocone (mesiolingual) are forming the trigon, while the hypocone (distolingual) is part of the talon. The rectangular m1 is divided into trigonid and talonid. The trigonid is characterized by paraconid (mesial), metaconid (mesiolingual), and protoconid (mesiobuccal). Hypoconid (distobuccal), entoconid (distolingual), and entoconulid (lingual) are forming the talonid. The rectangular m2 is also divided into trigonid and talonid. Protoconid (mesiobuccal) and metaconid (mesiolingual) are made up of the trigonid, while hypoconid (distobuccal), entoconid (distolingual), and entoconulid (lingual) are part of the talonid. The m3 is rounded and wrinkled with no defined cusps. In 23 specimens the metaconid (mesiobuccal) is present.

All 27 specimens are characterized by a bicuspid metaconid at the m1. The metaconid at the m2 is in 21 specimen bicuspid.

Nearly all of the analyzed specimens show the same cusp pattern. In one case there is an additional cusp mesial of the protocone at the P4 (SNSD B8309). The M1 of SNSD B21795 is characterized by an additional cusp between protocone and hypocone. The metaconid at the m2 of SNSD B16373 is made up of three instead of two cusps.

***Ursus maritimus* PHIPPS, 1774 – polar bear**

The triangular P4 is characterized by a prominent paracone (mesiobuccal), metacone (distobuccal), and a reduced protocone (lingual). The quadrate M1 is divided into trigon and talon. Paracone (mesiobuccal), metacone (distobuccal), and protocone (mesiolingual) form the trigon, and distolingually there is the hypocone distolingual). The M2 is nearly similar to the M1 except the elongated hypocone. The trigon consists of paracone (mesiobuccal), metacone (distobuccal), and protocone (mesiolingual). The rectangular m1 is divided into trigonid and talonid. The trigonid comprises the paraconid (mesial), metaconid (mesiobuccal), and protoconid (mesiobuccal). The talonid is made up of the hypoconid (distobuccal) and entoconid (distolingual). The m2 is also divided into trigonid and talonid. Protoconid (mesiobuccal) and metaconid (mesiolingual) form the trigonid, while hypoconid (distobuccal) and entoconid (distolingual) form the talonid. The m3 is rounded and wrinkled with no defined cusps. In 10 specimens the metaconid (mesiobuccal) is present.

Almost all of the analyzed specimens show the same cusp pattern. In three of 10 specimens the protocone at the P4 is lost. Only one specimen (SNSD B4550) shows an entoconulid at the m1 and m2, respectively. The metaconid of one specimen (SNSD B2428) is bicuspid.
